# Sensing rate, not size: growth rate-dependent negative feedback achieves the required organ size

**DOI:** 10.64898/2026.06.07.730717

**Authors:** Carmen M A Coelho, Abhishek Sharma, Varshitha Prabhakar, Huimiwon Ramwung Shimrah, Jaya Anusha Rachamadugu, Saher Chawla

## Abstract

The intrinsic mechanism of organ size determination remains elusive despite decades of research into the cellular nature, mechanics and genetics of growth in *Drosophila* wing imaginal discs and certain vertebrate organs. Using wing imaginal discs as a system, we propose a new model whose basis is growth rate dependent-negative feedback without organ size being sensed. We propose that such a model could correct for variations in growth trajectories and determine the average trend that each organ uses to reach the required size in a given time. We establish that alterations in size before the growth period are not corrected for, but alterations in growth rate during the start of the growth period induce size correcting feedback.

## Introduction

Despite a large number of studies on organ growth both in invertebrate and vertebrate systems, over the past two and a half decades, the mechanism by which final size is determined remains a mystery (for reviews see Hariharan, 2015); Penzo-Méndez and Stanger, 2015); Boulan and Léopold, 2021); Hernández-García et al., 2024). Work during the previous century suggested the possibility that size determination was influenced by the ability to somehow sense size (reviewed in Bryant and Simpson, 1984; (Day and Lawrence, 2000); (Coelho, 2020). In this study we explore the ability of growth rate itself to regulate growth rate, such that final size is determined without it being sensed.

Organ growth is regulated by both organ extrinsic and intrinsic signals (for reviews see Bryant and Simpson, 1984; (Boulan et al., 2015); (H’ng et al., 2023)) and an influence of extrinsic signals on intrinsic mechanisms (Parker and Struhl, 2020). Examining the essence of the problem, the size of an organ should be determined by the rate of its growth and the length of its growth period. However, growth rates are not constant during the growth period. After a period of quiescence, growth accelerates and then decelerates. The duration of these two phases could vary between organs. In vertebrate systems, using growth models fitted to weight and size data, it is substantially documented that growth rates are at their highest during the first one-third of the growth period ((Stewart and German, 1999); (Nijhout and German, 2012), after which there is a lengthened slow down. For the wing imaginal discs such detailed analysis is lacking, but a marked slowdown in disc size increase is detected during the later one-third of larval life (Bryant and Simpson, 1984; (Wartlick et al., 2011); (El Marzkioui et al., 2025); (Zhao et al., 2025). Wartlick, et al (2011) have described growth to be a process of continued slow-down from the 2^nd^ instar onwards, which comprises three fourths of the larval period. Is the phase of early deceleration or the phase of later slow down more critical in determining final size? It could be argued that the final slowing down phase is a key regulator of size as the organs are approaching their final size during this phase. Size sensing mechanisms could alter growth rate such that the appropriate final size is reached. Indeed, many hypothetical models have been proposed to explain how organ-size dependent patterning information and/or mechanical forces could alter growth rate during this final slowing down phase in wing imaginal discs ((Aegerter-Wilmsen et al., 2012); (Day and Lawrence, 2000); (Hamaratoglu et al., 2009); (Nijhout, 2003). However, these models remain unestablished.

Even if the final phase of slow-down is a critical determinant of final size, through a size sensing mechanism, how does this process of deceleration cope with a change in the initial rates of growth? Here we study the impact of early growth rates on the growth trajectory. We first describe the distribution of wing disc sizes during the larval growth period at two different temperatures, 25℃ and 29℃ and fit mathematical models to this data. We experimentally test the basic principles of these models and in doing so reveal the importance of studying the start of growth deceleration.

## Results

To understand how growth rate deceleration adjusts to changes in initial growth rates, we decided to study growth at two different temperatures. The larval period is 4 days at 25℃, but 3 and a half days at 29℃. Thus, larvae and the constituent organs are expected to initiate growth at a faster rate at 29℃. To describe growth trajectories at these two temperatures, wing disc size was examined from fixed samples at different times during larval life. We fitted polynomials to this data obtained from large cohorts of discs (see Figure 1A-B) and found that there was greater deviation from the polynomials in the first two-thirds of the period, than later in larval life (see Figure 1C-D) a phenomenon that is described statistically as heteroscedasticity. The basis for this deviation is analysed in Supplemental Information Sheets S1-S4. The analysis suggests that growth trajectories are not rigid, even under controlled culture conditions. One likely cause of variation is the slight differences in conditions between culture vials. However, another source of variation that is investigated below is the size differences that arise within paired organs of the same larva, due to stochasticity in the start of growth.

**Figure 1:**
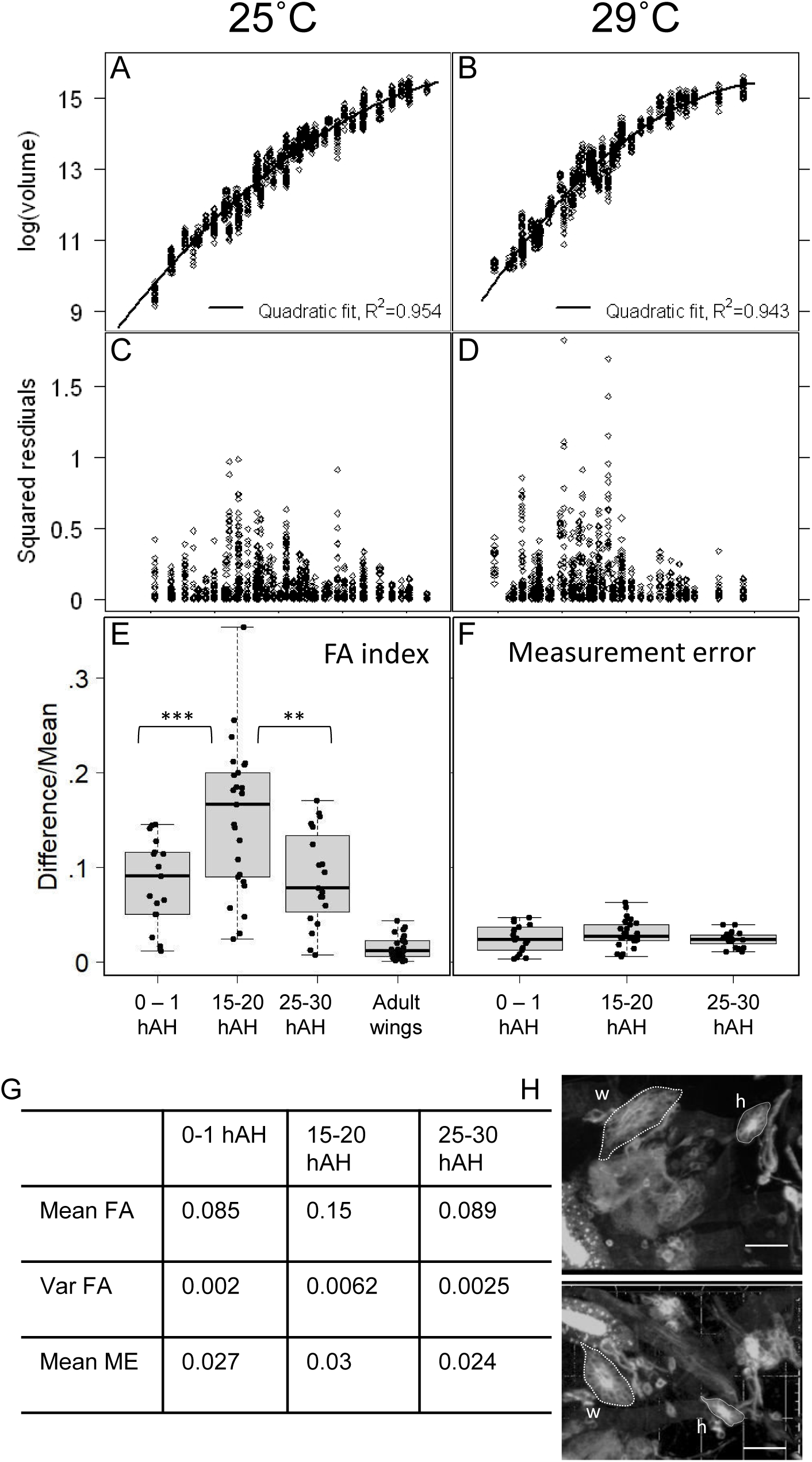
Analysis of variance in wing imaginal disc size during development. Wing discs were dissected from larvae at different developmental time points. The graphs here represent an analysis of variance in this data. Graphs **A,C** and **B,D** represent data from larvae cultured at 25℃ and 29℃ respectively. The x-axes of **A**, and **B** align with the x-axis of **C** and **D** respectively and show hours after larval hatching from the embryo, (hAH). Graphs **A** and **B** are semi-log scatter plots of log wing disc volume (Y-axis of **B** is aligned with that of **A**, n=1134 in **A** and 1021 in **B**), with quadratic polynomial fits (black lines) log Volume with hAH; the squares of the residuals from these fits are shown in **C** and **D** (Y axis of **D** is aligned with that of **C**). **E-H** represents the analysis of FA in wing discs during early development. Z-stack images were taken of the left and right disc of individual esg-Gal4;UAS-mCD8 larvae early in development and the larvae were placed back in the single larvae culture capsules to allow them to develop to adulthood enabling FA analysis in adult wing size. Each larva was imaged once, either within 1hr after hatching, between 15-20hAH or between 25-30hAH. FA index is calculated for each disc as absolute difference/mean of the pair and the spread of FA indices for each time interval is shown as box and whisker plots in **E**, with the statistical analysis tabled in **G**; the adult wing data in **E** is that of the larvae that survived the imaging process. The mean FA and variance in FA of the 15-20Hr group (see **E**) differs significantly different from the 0-1Hr group (p=0.001157 for mean (Welch two sample t-test) and 0.024 for variance (Fischer test)). The mean of the 15-20 Hr group varies significantly from the mean of the 25-30 Hr group (p=0.0023), but the variance does not differ significantly (p=0.056). Measurement error is measured as the difference in values obtained from two independent analyses of the same disc, divided by the mean of the two values (see Supplemental Information sheet S1) displayed in D reveals that the measurement error remained the same across sample groups. The images in **H** display the wing (w) and haltere (h) imaginal discs, with their boundaries marked by dashed lines, in the right (image above) and left (image below) sides of a larva showing high FA in wing disc volume; scale bar represents 30µm.

### Developmental Stochasticity

Soon after hatching from the egg, larvae start feeding and this triggers an exit from quiescence in larval tissue ((Galloni and Edgar, 1999)). The exit from quiescence in imaginal tissue has been poorly studied. The only study of growth of the imaginal discs during these early stages is the histological analysis by Madhavan and Schneiderman (1977), in which they described a six-fold increase in cell volume within the first 15 hrs of larval life, in most of the imaginal discs ((Mandaravally Madhavan and Schneiderman, 1977)). Histological studies have their limitations. These studies did not reveal the actual start of growth and the small numbers did not allow an analysis of stochasticity.

Since it is difficult to dissect apart imaginal discs during the first instar stages, we examined disc size in larvae expressing GFP in imaginal tissue in a strain that will henceforth be referred to as esg-Gal4-GFP (see materials and methods and Supplemental Information). As shown in Supplemental Table S1, mean disc size at time points ranging from 0 to 6 h after hatching (hAH) remain invariant, consistent with a lag during this period. However, one sample set (larvae from one culture vial) at 6hAH, showed a much larger volume, indicating that growth of the wing discs has begun in this set of larvae, indicating that the start of growth of the wing imaginal discs is at around 6hAH. The variability between vials could be attributed to slight differences in culture conditions amongst them. To detect developmental stochasticity, we need to analyse variation between left and right discs after the start of growth.

To examine if there was indeed stochasticity in the start of growth, we examined differences in the size of the right and left wing-disc in larvae at three different time points: 0 to 1, 15 to 20 and 25 to 30 hAH in esg-Gal4-GFP larvae. We did this in live larvae and so were able to put the larvae back inside food in single larva culture vials, after taking the images of the left and right wing-discs, enabling an analysis of FA in the adult wings in the same individuals. We used Desflurane to immobilise larvae during image acquisition. We have found that repeated treatment does cause some effect on development and reduces survival. Hence, larvae were imaged at just one of the above time points and placed back in the food. A box and whisker plot of the FA indices for each sample set shown in Figure 1E shows that FA exists even in newly hatched larvae. FA index is calculated for each disc pair as the absolute difference divided by the mean volume of the pair (see (Palmer and Strobeck, 1986) for methods of calculating FA index). This FA increases significantly in the 15 to 20 hr sample and then decreases again in the 25 to 30 hr sample. The 15-20hAH sample shows an increase in both the mean FA and the variance in FA (see Table in Figure 1G), suggesting that there is an amplification of size differences as discs start to grow. This amplification of size differences can only be due to differences in the time of the start and/or the differences in the rate of growth at start. A similar pattern of FA during early larval life was observed through the analyses of fixed samples (data not shown). As revealed in the box and whisker plot shown in Figure 1F and the Table in Figure 1G, the average measurement error is lower than the average FA index and does not vary with age of the discs (see Supplemental Information for analysis of measurement error). Thus, this analysis does reveal that FA increases during the first instar of larval life, and starts to decrease within approximately 10 hours. The FA at 15-20 hAH is indicative of developmental stochasticity in the start of growth, with the beginning of a correction evident 10 hours later. Adult wing FA of the survivors shows that a correction in early FA has certainly taken place.

### A model to explain the decrease in variance

Whatever may be the cause of the variance in disc size during larval development, the decrease in variance at the end would require a correction in growth trajectories. What causes this correction? It would be reasonable to expect that faster growing discs slow down their growth faster and/or at an earlier stage in development than slower growing discs. And conversely, slower growing discs would undergo a slower deceleration in growth and/or a later deceleration in growth. This implies that growth rates correlate not with disc size or developmental stage, but with “history”, or the trend in previous growth of the disc.

To explore this possibility, we required a model in which size does not feedback onto growth rate and there is scope for an accumulating form of feedback, that would provide a plausible mechanism for the effect of history on the growth trajectory. To develop such a model, we first used matlab to fit a set of Ordinary Differential Equations (ODEs) to our volume data.

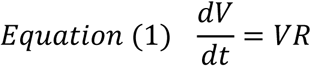

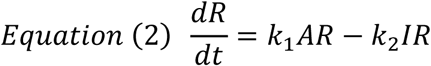

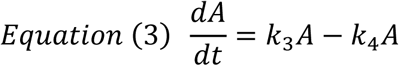

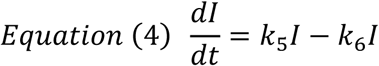

From Equation (1), we derive a definition for R.

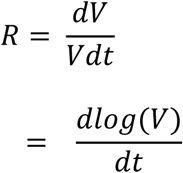

Closed form solutions for these equations can be obtained. However, we solved the equations numerically by trying different starting values for the initial conditions and parameter values. We ran several simulations using Matlab and chose the ones that best fit the experimental data based on the correlation coefficient, R^2^. Some simulations gave negative values for the ‘k’ constants. These were not taken except for one model (see Supplemental Table S2), where both k3 and k4 had negative values and so in effect replaced each other.

We found initial conditions and values for the constants k1 to k6, that enabled good fits to the 25℃ data and the 29℃ data with R^2^ values = 0.945 and 0.954 respectively (see black and grey curves in Figure 2A; black is fitted to the 29℃ data and grey is fitted to the 25℃ data). The constants and initial values obtained for this model and for the models below are tabulated in Supplemental Information Table S2. In this model and the models below, R, the rate of growth acts on volume (V), but V does not act back on R. R changes with time and is dependent on the levels of activator (A) and inhibitor (I) and the constants k1 and k2, as represented by equation 2 (see Figure 2E). I and A themselves change with time dependent on their rates of synthesis and degradation as represented in equations 3 and 4 (see Figure 2I and M). Since the experimental data does not include the first quarter of the larval stage, namely the first instar, we used these initial conditions and constants, to simulate backwards (see dotted lines in Figures 2A, E, I and M) and thus obtain the initial values at the start of larval life, corresponding to discs within larvae freshly hatched out of the egg.

**Figure 2:**
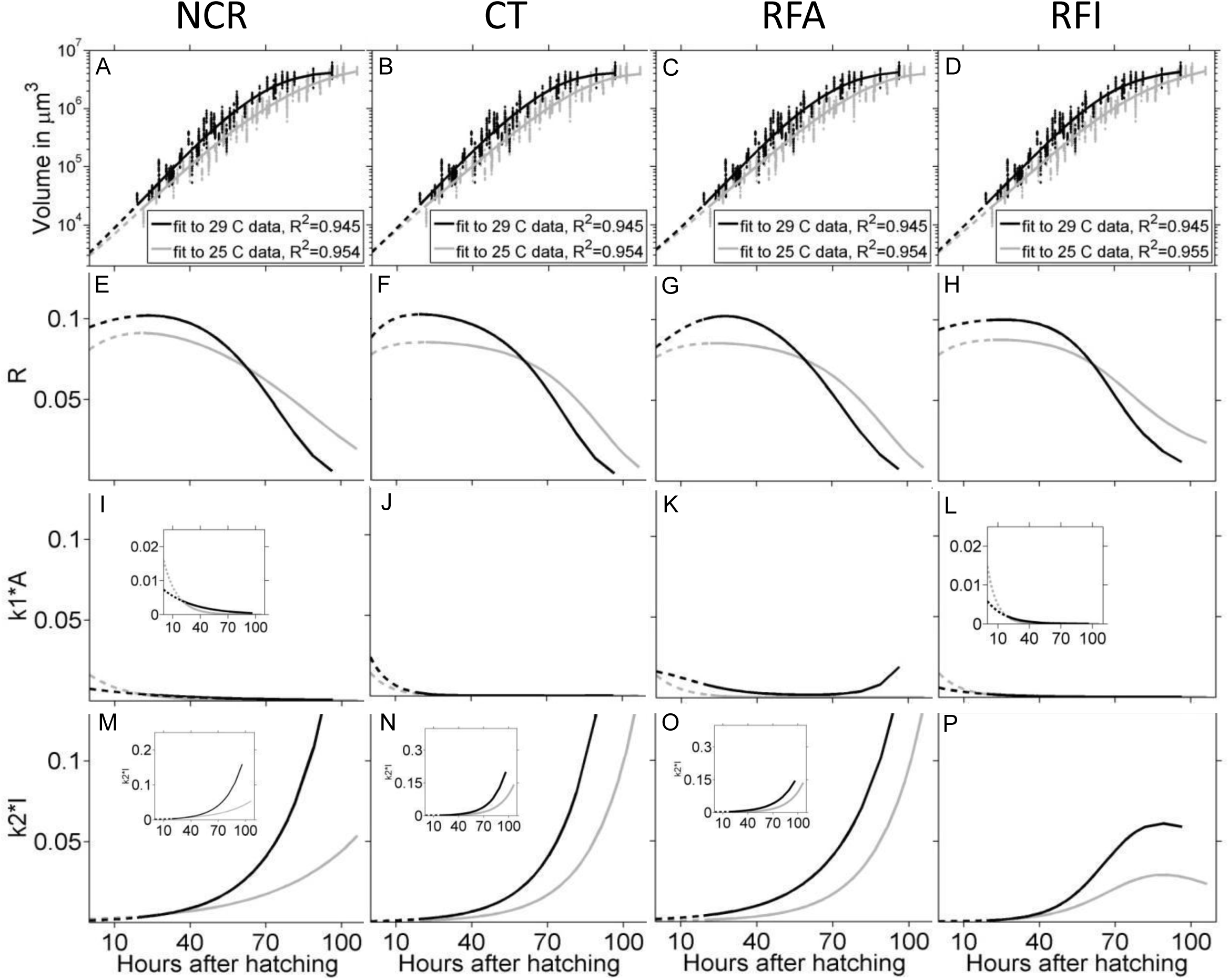
Fits of different mathematical models to wing disc volume data versus developmental time. Mathematical models NCR, CT, RFA and RFI (see text for expansion of these names) were fit to volume versus time. A-D are semi-logarithmic plots of volume versus time, with black and grey dots representing wing disc volume from larvae cultured at 29℃ and 25℃ respectively. In all graphs A-P, black and grey curves represent fits to 29℃ and 25℃ data respectively. The graphs are arranged such that one column represents one model (with the name of the model indicated above the column) and the x axes for each column align with that of graphs in the bottom most row, M, N, O and P respectively. Curves in A-D represent the volume fits with the Y-axes aligning with that of graph E. These volume fits are a result of R acting on V. The change in R with time is shown in graphs E-H, with the Y-axes aligning with that of E. This change in R is brought about by the sum of k1*A and - k2*I acting on R at every time point. The changes in k1*A and k2*I with time are shown in graphs I-L and M-P with Y-axes aligning with graphs I and M, respectively. In all models the k2*I values rise much higher than the k1*A values and the rise in k2*I is lower in the RFI model as depicted in graphs in P. To demonstrate these differences, the scales are identical for the k2*I and k1*A graphs, with detail shown in the insets in I and L. Since the k2*I values rise higher than these axes in M, N and O, insets are used to show the entire graphs. In all graphs the dotted lines represent the values obtained through backward simulation (see text).

To test if this model can explain how initial variation in disc size could be reined in, we introduced random variation in the initial conditions, and simulated growth until the end of larval life. Graphs in Figure 3A and E shows twenty such simulations using the initial values obtained from the 29℃ fits, when R_0_ is varied and graphs in Figure 3I, M and Q shows twenty simulations when A_0_ is varied. Simulations in which I_0_ is varied are shown in Supplemental Figure S6 (see Supplemental Figure S6 A, E, I and M). Similar results were obtained with the 25℃ fits (data not shown). It is clear that in all these simulations the differences in the volume trajectories become more pronounced with time. Therefore, this model is not able to bring about a decrease in size variation towards the end of the growth period. We named this model “No cross regulation” or NCR. To see if feedback could make the volume trajectories converge over time, we fit models that included feedback components. Three such models were tested:

**Figure 3:**
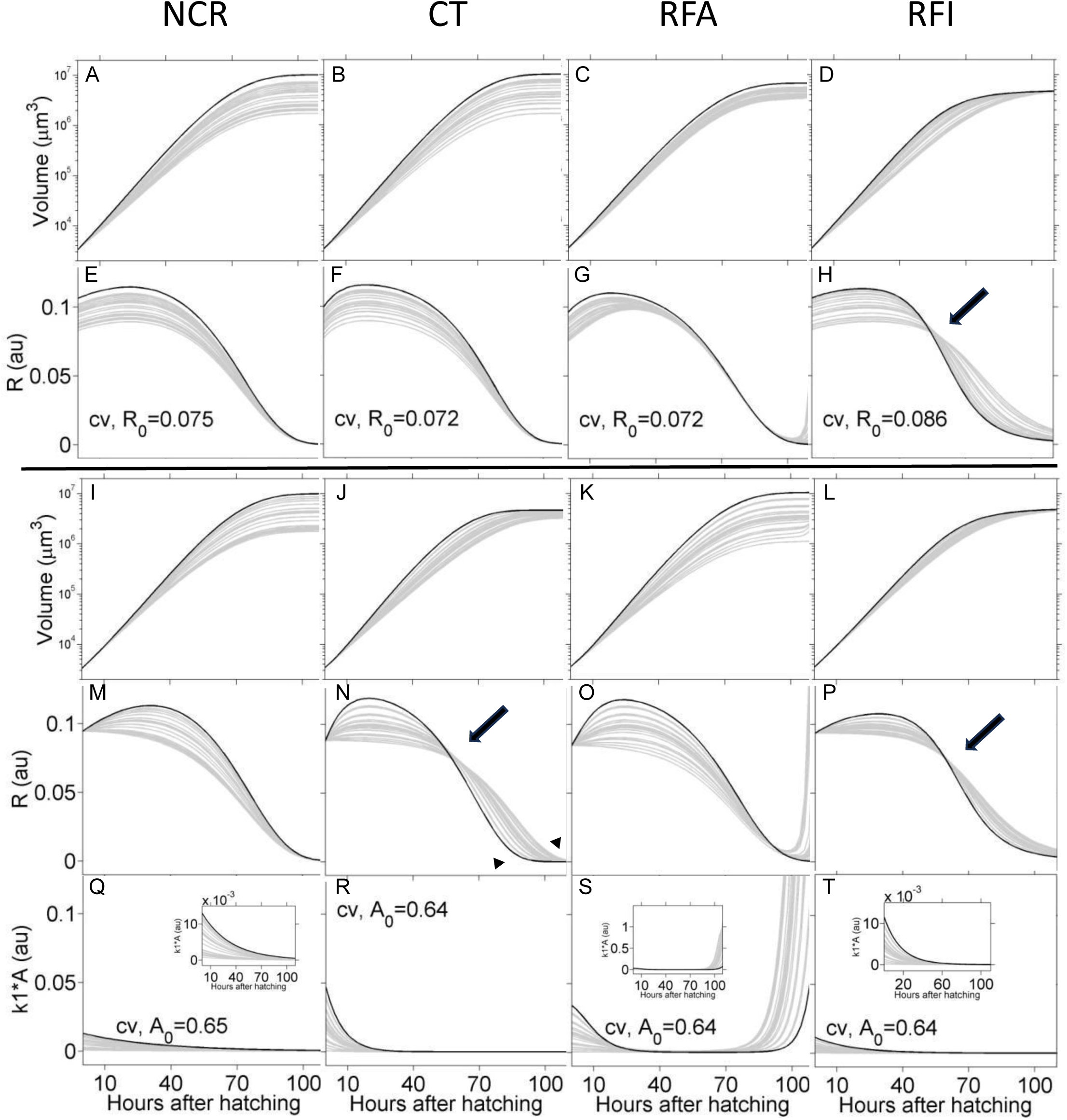
Simulations to test which model allows for correction of growth trajectories. Random variation was introduced in the initial values for either R (graphs A-H) or A (graphs I-T). The initial values, representing the start of the growth period were obtained from the backward simulations depicted in Figure 2 (dashed lines in Figure 2, see text). The names of the mathematical models are as depicted above each column (see text for expansion of the names). The curves representing the highest initial value of R_0_ are depicted as a black curve. The x-axes for each column align with that of the x-axis of the graph in the bottommost row (Q-T). The Y-axes for graphs in each row align with the Y-axis of the graph on the left. Variation in R_0_ induces variation in the R curves (**E-H**) and the V curves (**A-D**) for all models. Variation in k1*A and k2*I is induced with the RFA and RFI models respectively and is shown in Supplemental Figure S4. Variation in A_0_ induces variation in the A (**Q-T**), R (**M-N**) and V (**I-L**) curves for all models. Variation in k2*I is induced only with the CT and RFI models and is shown in Supplemental Figure S5. The variation in the initial values was introduced using a random number generator, ensuring that the coefficient of variance (cv) was comparable for all models (R_0_ at approximately 0.072 and A_0_ at approximately 0.064). Larger cv was used for R in the RFI model (H) as the volume curves converge more effectively in this model. The Y axes scales are the same as that used in Figure 2, for ease of comparison. Where necessary, details are shown in the insets. Note that the R curves cross (arrow) in approximately the middle of the simulation period in graphs shown in H, N and P. and the black curves reach the lowest values at the end of simulation period in the RFI model column. In the corresponding volume graphs (D, J and L), the variation in volume is visible in the middle of the simulation period, but decreases at the end. In all other volume graphs, variation in volume does not reduce at the end of the simulation period.

Model 1 is called “Rate dependent feedback on Activator” or RFA. In this model, an additional term is introduced into equation (3), such that the rate of degradation of the activator A is influenced by the rate of growth R.

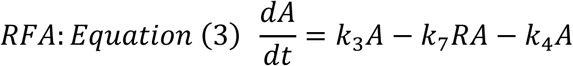

Model 2 is called “Rate dependent feedback on Inhibitor” or RFI. In this model an additional term is included into equation (4), such that rate of synthesis of the inhibitor I is positively influenced by the rate of growth R. A model involving the synthesis of inhibitors was proposed by Nijhout ((Nijhout, 2003). In his model, the cumulative effect of the inhibitor on growth is a result of the increase in the number of cells that synthesize the inhibitors, the number of cells serving as a proxy for the size of the organ. In the model described in this study, inhibitor levels depend on the rate of growth R, which determines growth rate locally and is not influenced by the size of the organ as explained above.

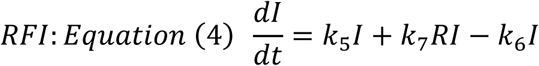

Model 3 is called “Cross talk” or CT. In this model an additional term is included into both equations (3) and (4), such that the levels of the inhibitor influence the rate of degradation of the activator and simultaneously levels of the activator influence the synthesis of the inhibitor.

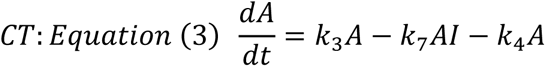

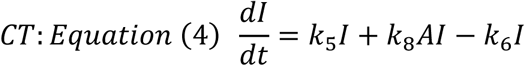

We fit these models to the experimental data and obtained the initial values and values for the constants (see black and grey curves in Figure 2). We then used the respective constants and initial values to simulate backwards (see dotted lines in Figure 2) to obtain the initial values that would correspond to the start of larval life, as was done with the model NCR. We then introduced random variation in the initial values and ran simulations to see which model allowed a convergence in final volume. In Figure 3, we depict the results obtained from the 29℃ fits. Similar results were obtained from the 25℃ fits (data not shown).

The changes in V and R as a result of the simulations are depicted graphically in Figure 3 (see Figure 3A-H for simulations resulting from variation in the initial values of R and see Figure 3I-T for simulations resulting from variations in the initial values of A). The trajectories with the highest initial value of R, are depicted in black, while other trajectories are in grey. As can be seen only RFI is able to bring about a convergence in size trajectories at the end of the simulation, when random variability is introduced in R_0_ (see curves in Figure 3D). As seen in Figure 3H, the R curves cross each other around the middle of the simulation period (see arrow in Figure 3H). High initial R values result in steeper and earlier deceleration and lower R values at the end of the simulation. The black curve representing the highest R_0_ value ends with the lowest R value at the end of the simulation period. On the other hand, the R trajectories converge towards the end of the simulations in NCR, RFA and CT, without crossing each other. R increases at the end of the simulation in RFA.

When A_0_ is varied at the start (see Figure 3 I-T), both CT and RFI are able to bring about convergence in volume at the end of the simulations (see Figure 3J, L). In both these models, R values result in steeper and earlier deceleration, with the R curves crossing midway during the simulation period (see arrows in Figures 3N and P). With the RFI model, the trajectories that start with the higher R values end with the lower R values at the end of the simulation (see black curve in Figure 3P). Here again, R and A increase at the end of the simulation in RFA with no convergence in V.

When I_0_ is varied at the start, none of the models are able to bring about a convergence in size at the end of the simulations (see Supplemental Figure S6). Since RFI is able to bring about convergence in end volume when either R or A are varied at the start, this was considered to be the most robust model. Simulations were performed using the RFI model, suitably varying together, V_0_, R_0_, A_0_ and I_0_ such that a set of 20 simulations was able to mimic the variance observed in the experimental data at both 25℃ and 29℃ (data not shown). A similar exercise using the CT model was able to mimic the 29℃ data, but not the 25℃ data (data not shown). In summary, the fits of these models to the experimental data and the simulations reveal that for the correction of variance in size trajectories, faster growing organs need to slow their growth earlier and faster than the slower growing counterparts, such that the R curves cross each other. RFI is a more robust model than CT. It should be noted that in all models the peak rates of growth are during the first quarter of the growth period (see Figure 3M-P).

### Fitting the RFI model to larval growth trajectories

This far we have examined variance in disc size using data pooled from multiple experiments. The growth trajectories of individual discs have not been followed through the entire length of the growth period. Since it is not possible to do so for the entire growth period with the required resolution in time intervals for the wing imaginal discs, we followed the growth trajectories of individual genetically isogenized CS larvae. Single larvae were collected soon after hatching from synchronized egg lays and cultured individually in single larva culture capsules and maintained at 25 ℃ and 29℃. The volume of the larvae was calculated from measurements of the heights and widths of the larvae, from bright field photographs taken every four hours, until pupation. Only female larvae were considered in the final analysis. The 25℃ data is depicted in Figure 4 and the 29℃ data is depicted in Supplemental Figure S7. As can be seen from the graphs of log volume against developmental time (scatter plot in Figure 4A), there is some spread in the volume data during development, but the spread is reduced by the end of larval life. The RFI model was fit to the collective data to obtain average initial values and the values for the constants k1 to k8 (shown in Supplemental Table S3). Keeping the constants thus obtained, as fixed values, the model was fit to the trajectory of each larva separately. The V, R and I curves thus obtained, are depicted in Figure 4 B-E. They are colour coded according to starting R values. The derivatives of the R curves reveal the change in the rate at any point in development. When these values are positive, growth is accelerating and when these values are negative, growth is decelerating. The accelerating phase is brief for all curves as seen in Figure 4D. Thus, growth starts to decelerate soon after the start.

**Figure 4:**
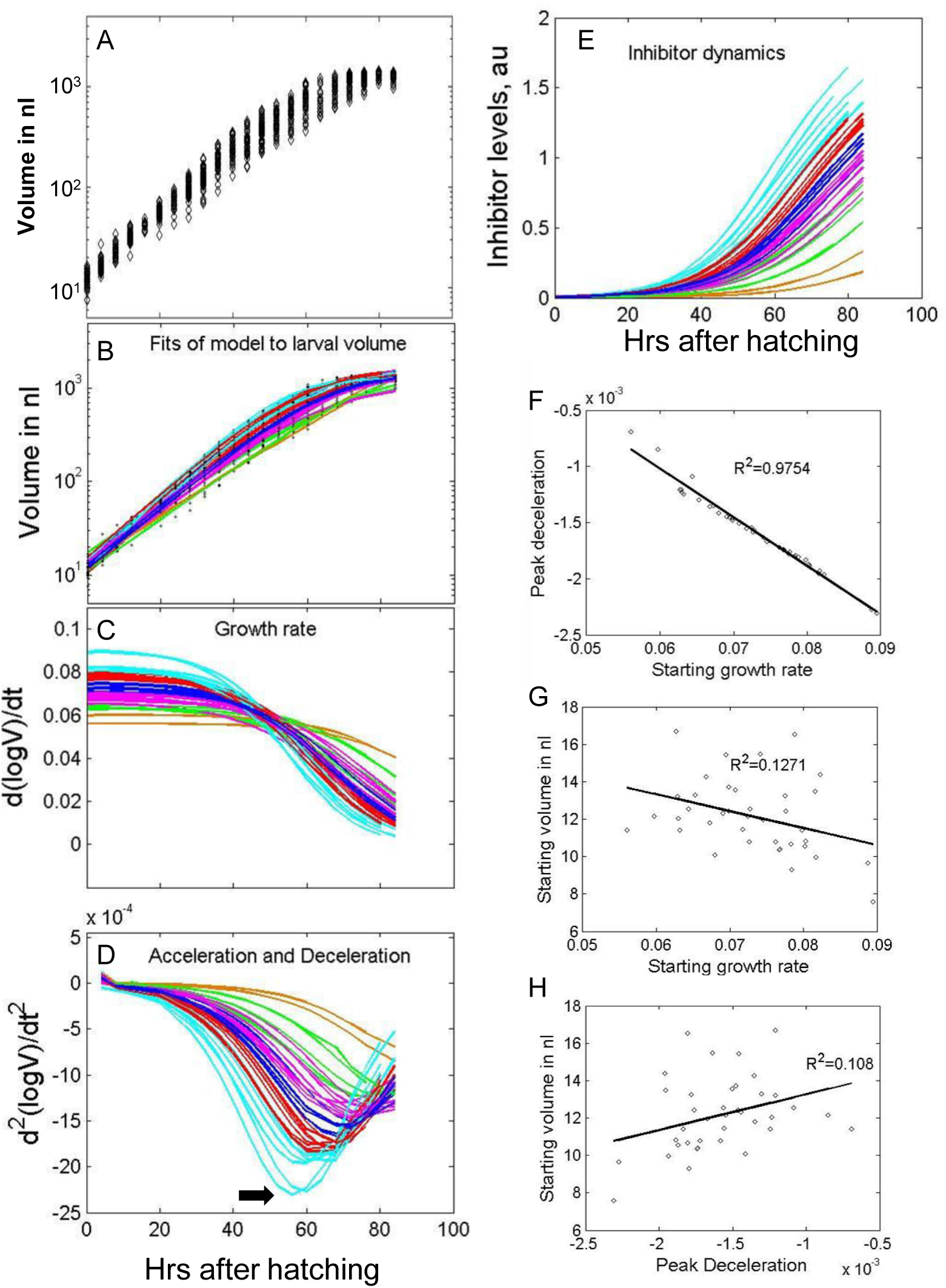
Growth trajectories of isogenized larvae cultured individually. Isogenized larvae were cultured in single larvae culture vials at 25℃, they were immobilized every four hours and volume was estimated from the bright field images taken (see materials and methods), shown here in A as a semi-logarithmic plot of volume versus hours after hatching (hAH). The RFI mathematical model was fit to this data and is shown in graphs B (volume), C (R) and E (I). The graphs are color coded according to the starting value of R (Cyan, red, blue, magenta, green and yellow ochre; highest to lowest initial R values). D represents the derivatives of the curves shown in C. The curves reach their minima and then start to increase again. The minimum of each curve is referred to as the peak deceleration value (PDV, arrow in D). Charts F to H represent regression analysis between starting growth rate, starting volume and PDV, with the value of R^2^, the Regression coefficient indicated. The x-axes of graphs A-D align with that of D.

One can see that the V curves cris cross and are not parallel to each other. The R curves also criss-cross and the curves with the highest starting R values (cyan curves) show the steepest declines and end with the lowest values (see graphs in Figure 4C). The derivatives of the R curves also reveal that the larvae with the fastest initial growth rates (cyan curves) undergo the steepest deceleration, reaching a minimum value (referred to henceforth as the peak deceleration value (PDV)) before the curves rise towards decreasing negative values, towards the end of the growth period. The PDV is reached at earlier time points for the larvae that start with the fastest growth rate. On the other hand, the larvae that start with the lowest growth rates (ochre yellow graphs) maintain their growth rate for most of the growth period (see ochre yellow R curves in Figure 4C) and the deceleration values remain close to zero during this period (see corresponding ochre yellow curves in Figure 4D). Correspondingly, the cyan inhibitor levels show the steepest rise and highest levels at the end of the growth period and the inhibitor levels for the ochre curves show the slowest and most shallow increase (Figure 4E).

There is some variation in the starting volumes of the larvae. To examine whether the larvae with the smaller volumes are the ones that undergo the fastest starting growth rates, a regression analysis was performed comparing starting volume with starting R and starting volume with PDV and starting R with PDV. As can be observed in Figures 5F-H, there is no correlation between the starting volume and starting R and also starting volume and PDV. There is a very strong correlation between starting R and PDV (Figure 4F). Regression analysis between other values reveals that there is a strong but weaker correlation between starting R and time at which PDV is reached (see Supplemental Table 4). Thus, this analysis reveals the influence of starting R values on the rest of the trajectory and the lack of influence of starting volume on the rest of the trajectory.

**Figure 5:**
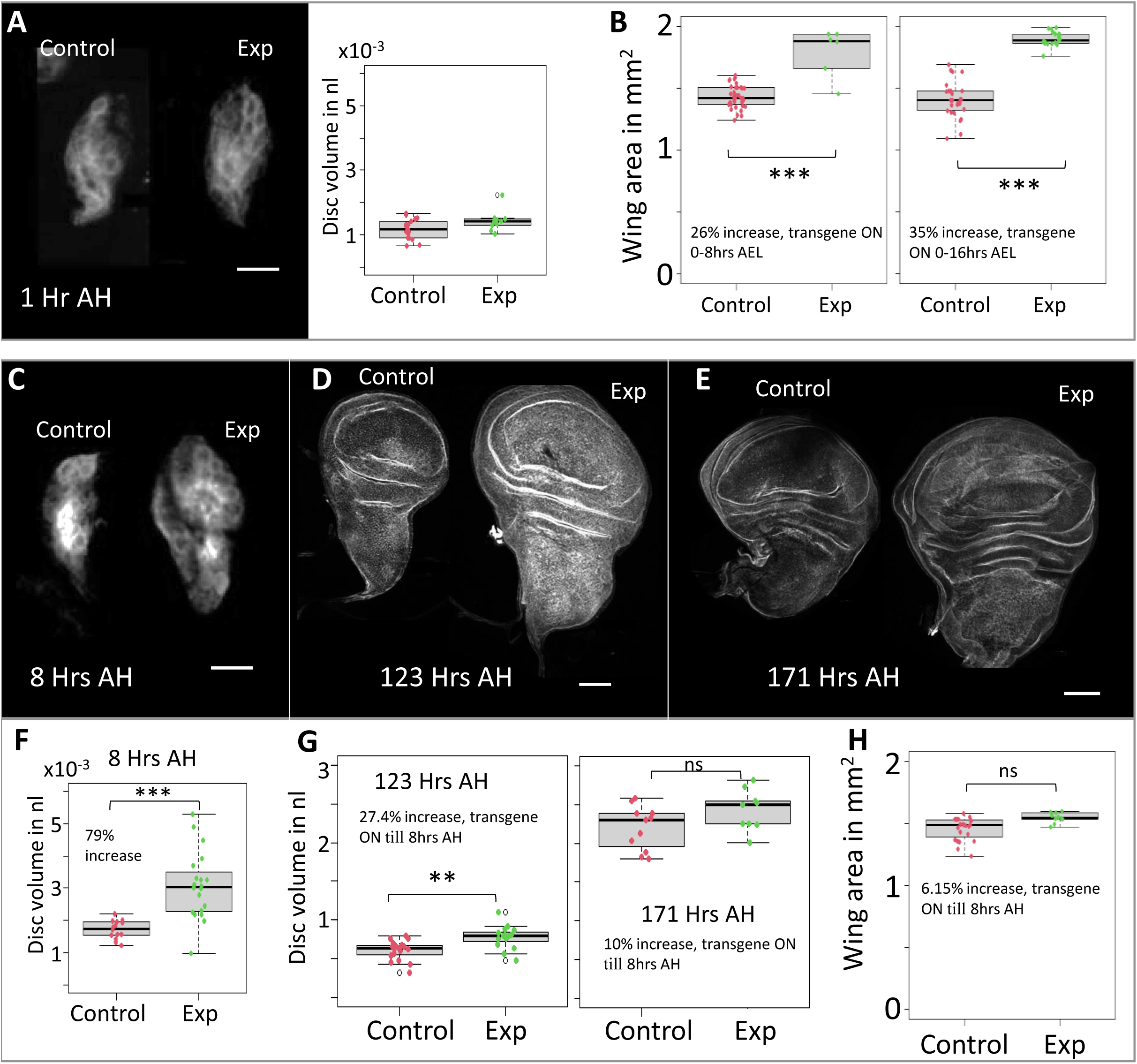
Starting the growth period at a bigger size versus at a faster growth rate. The temperature sensitive Gal80^ts^/Gal4 system was used to drive expression of an activated form of class1A PI3Kinase, Dp110 (Dp110-CAAX, Leevers, et al, 1996) during the embryonic period (A-B), or during the embryonic period and into the first 8 hours of larval life (C-H). Expression of Dp110-CAAX during the embryonic period results in larger sized adult wings (B) in Experimental (Exp) versus the only driver control (Control). The period of transgene expression (period at 29℃) is as indicated in each graph along with the % increase in mean wing area in experiment versus control; AEL implies time <u>a</u>fter <u>e</u>gg lay. Images of such discs could not be taken as the expression of GFP fades when the temperature of Gal80 is brought down to 18℃, its permissive temperature. The charts and images in A serve to illustrate the increase in size that occurs due to such expression until 1 hAH. On the other hand, if expression of Dp110-CAAX was continued into the start of the growth period, 8 hAH (images of discs in C-E and box plots in F-H), the 79% increase in mean size was brought down to a 27.4% increase at 123hAH, 10% increase at 171 hAH and 6.15% increase in mean adult wing area, the latter two being statistically non-significant. Scale bar in A and C represents 10µm, in D 50µm and E 100µm.

### Results of altering volume before the start of growth

In the models proposed above, the size of the organ or animal does not feedback onto growth rate. If this were indeed true, an alteration in size before the start of the growth period would not be corrected for and these animals would retain the same size alteration until the end of the growth period. On the other hand, an alteration in rate of growth during the growth period could initiate rate correcting feedback. To test these predictions, we used the Gal4/Gal80^ts^ system to temporally regulate the expression of a growth promoter, constitutively active class 1A PI3Kinase, membrane targeted Dp110 (henceforth referred to as Dp110_CAAX, (Leevers et al., 1996)) during different developmental stages. *escargot-Gal4* was used to drive transgene expression and temperature sensitive Gal80 was used to temporally modulate the expression, with the Gal4 being active at 29℃ but inhibited by Gal80^ts^ at 18℃.

The embryonic period of *Drosophila* is a period of no growth. Indeed, we have observed that interfering with growth regulating pathways during the embryonic period with the expression of a kinase dead version of the PI3Kinase, Dp110 does not reduce disc size (data not shown). But, driving Dp110_CAAX during the embryonic phase of development until 1hAH, results in increased disc size. Mean disc volume of Dp110_CAAX discs is 22.2% larger in the controls (see box and whisker plot in Figure 5A). As mentioned above, there is an approximately 6h time lag before growth starts at 25℃. At 18℃ this time lag could be longer. The eggs were allowed to develop at 29℃, at which time Dp110_CAAX is expressed and then transferred to 18℃, when expression of the transgene would be shut down and growth would be regulated by the endogenous wild type Dp110.

At 29℃, the embryonic period lasts for 17½ hours. Since we did not have an estimation of the turn-over rates of the transgene upon expression, the eggs were kept at 29℃ from the time of egg lay to 9 hours before hatching and 1 hour before hatching (0 to 8 and 0 to 16 h after egg lay (hAEL), respectively). This would allow sufficient time for the degradation of the induced Dp110_CAAX before the start of growth. If the induced Dp110 increased disc size, these samples would represent a situation where size is increased before the start, but rate of growth at start is not altered. On the other hand, when the cultures are maintained at 29℃ until 8 hAH, a 79% increase is seen in mean volume of Dp110_CAAX discs compared to control (see images in Figure 5C and corresponding box and whisker plots in Figure 5F). These discs are larger than control and experience a higher-than-normal rate of growth during the start of the growth period.

As seen in Figure 5B, larger size before the start resulted in an increase in the mean area of the adult wings by 26% and 35% when compared to controls (see box and whisker plots in Figure 5B). However, larger size and rate during the start, results in just a 6.15% increase compared to control in adult wing size and this increase is not statistically significant (see box and whisker plot in Figure 5H).

We looked at disc size in wandering larvae which were maintained at 29℃ until 8hAH. Both experimental and control larvae were found to wander at similar times after hatching. But, a proportion of larvae wander earlier than others. When wandering larvae are dissected at 123hAH, the difference in mean volume is 27.4% (see images in Figure 5D and box and whisker plot on the left in Figure 5G) and when dissected at 171hAH, the difference is just 10% and is not statistically significant (see images in Figure 5E and box and whisker plot on the right in Figure 5G).

These results together suggest that when size is increased before the start of the growth period, this altered size is not corrected for. However, when growth rate is increased at the start of the growth period, a correction is initiated.

### What happens at the end of the growth period?

This far we have shown a strong impact of initial growth rates on the growth trajectories of both wing imaginal discs and whole larvae. We have also shown that alterations in volume before the growth period are not corrected for. If the normal growth period was extended, would discs stop growing at a particular size? In *Drosophila,* the length of the growth period is determined organ-extrinsically, by systemic circulating hormones secreted in pulses by the neuroendocrine system (reviewed in (Nijhout, 2003); (Nijhout et al., 2014); (Boulan et al., 2015)), without input from the developing organs during normal development ((Stieper et al., 2008); reviewed in Coelho, 2020). When flies are subjected to a day night rhythm, the time of larval wandering becomes influenced by the photoperiod ((Roberts et al., 1987)). We subjected fly cultures for five successive generations to 24 hr light and dark cycles; 16hr light and 8 hour dark phases, approximately in synch with the natural day-light rhythm. After five successive generations, we seeded freshly hatched synchronized larvae at fixed densities into vials just before the start of a dark phase and just after. Larvae that were seeded into vials before the start of the dark phase, wandered at the start of the 5^th^ light phase, while larvae that were seeded after the dark phase wandered at the start of the 4^th^ light phase, resulting in two different growth periods, 98 hours and 109 hours (see schematic in Figure 6A). We used two different food strengths (25% and 100%), to allow for the possibility that larvae that had eaten richer food may undergo faster deceleration, resulting in very slow growth rates towards the end of the growth period. Under such conditions a lengthening of the growth period would not result in a significant final size increase. However, in both food conditions we found that the discs from larvae that had undergone a longer feeding period, were bigger (see images in Figure 6B-C and graphs in Figure 6D). Thus, a longer growth period allows discs to reach a larger size.

**Figure 6:**
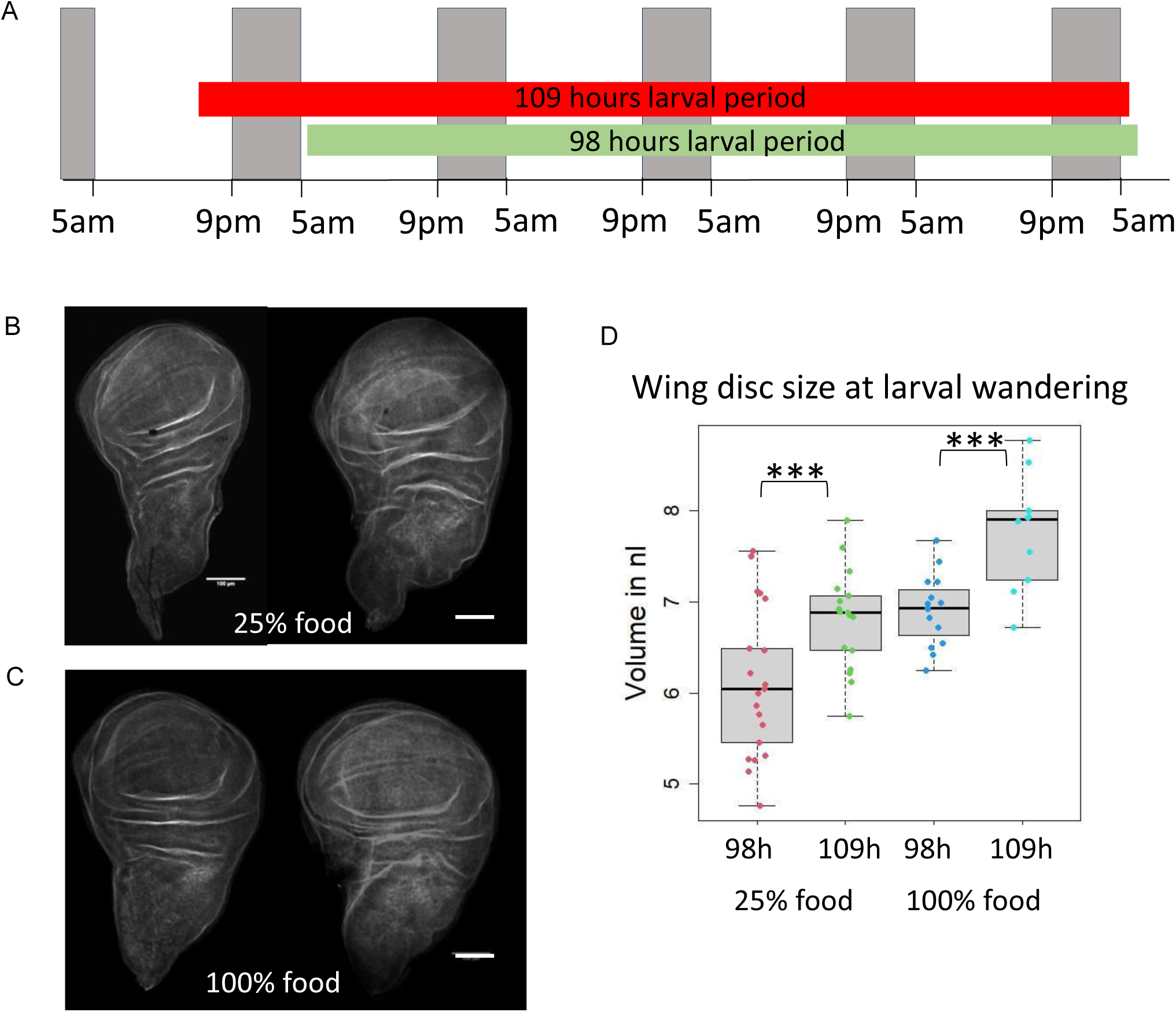
Increasing the length of the growth period allows increase in final disc size. Canton S flies were subjected to five generations of a dark light cycle and then newly hatched larvae were collected from synchronized egg lays just before (start of the red bar in A) and just after the dark cycle (start of the green bar in A) and cultured in food vials at fixed densities, one batch of larvae in 100% food and one batch in 25% food (see materials and methods). The ends of the red and green bars depict when the larvae wandered out of the food and were dissected. Images in B and C represent the mean sized discs; disc on left indicating the 98-hour larval period and those on the right depicting the 109-hour larval period. The box plots in D reveal the range of volume measurements. In both types of food, the volume of discs that underwent a longer larval period were significantly larger (p=0.0019 and 0.00058 in 25% and 100% food respectively). Scale bar in images B and C is 100 µm.

## Discussion

Studies this far on size determination of the wing imaginal discs have focussed on the later stages during which discs approach their final size. This is the first study analysing the impact of the start of growth on the rest of the trajectory. Growth of most organs start from a phase of no growth or quiescence. To reach a particular size within the available developmental time, they experience a phase of acceleration and then deceleration, the rates and duration of each help in determining when they approach their final size. Making use of the ectothermic nature of insect physiology, we started with the question of how temperature determined initial rates of acceleration would drive rates of deceleration. Because we studied a large sample size, we revealed heteroscedasticity of disc size when plotted against developmental time, even within a temperature. This observation led us to show that there is stochasticity in the start of growth. To correct this stochasticity, we propose that the initial rate of deceleration drives the rest of the trajectory, without the need to sense size, forming the basis of a new and novel model for organ size determination.

### Features of the models and the dynamics of deceleration

The models presented here are abstract in nature and do not imply any identified molecule or molecules. They utilize hypothetical activators and inhibitors that are present throughout the organ, but act to regulate growth locally, without any sense of position within the organ. The activators kick-start growth. In the models presented here, the activator levels decrease with time. However, we have also been able to fit these models with the activator levels remaining constant (data not shown).

Although we have been able to fit four different models to our experimental data, only two of these models reveal an ability to correct size variation when variation is introduced in the initial values of either A or R or both (see Figure 3). These models are the “Cross talk” (CT) model and the “Rate dependent feedback on inhibitor” (RFI) model. The CT model is not able to reduce size variation caused by variation in R_0_. This is not surprising because CT involves cross talk between A and I with no feedback from R in any of the equations. The RFA model invokes growth rate dependent degradation of activator. One reason why this model does not bring about disc size convergence is that the activator levels start to increase once rate drops to the low level that is characteristic of the end of the growth period. The parameters of A, R and I at this stage are not sufficient to keep A and R down. This doesn’t mean that rate dependent degradation of activator is not a biologically conceivable model. The set of ordinary differential equations used in this study do not provide a workable framework to test this mechanism.

Examining the behaviour of the R curves provides experimentally testable insight into what brings about disc size convergence. As growth slows down towards a near halt, differences in R values between growth curves during normal growth can be expected to decrease if not diminish completely. An interesting observation we make is that if varying volume trajectories are to converge, the R values should not converge at the end. In all simulations where the R curves converge, V curves in fact become parallel to each other and differences in V that manifest during the middle of the simulation get retained. The differences do not decrease. Convergence in disc size is accompanied by two R curve behaviours. One is that the R curves cross each other in the middle of the simulation. The other behaviour is that the R values remain different even at the end of the simulation. In the CT model, disc size convergence is accompanied by R values equalising as the end of the simulation approaches (see arrowheads in Figure 3N). Interestingly we observe that when the RFI model is fit to individual larval trajectories, the R values of each trajectory are different at the end of larval life. It can be experimentally tested whether differences in protein translation rates between organs and between larvae at the end of larval life correlate negatively with differences in starting growth rates.

### The mechanism for growth-rate dependent negative feedback

Our experimental data strongly suggests that the negative feedback is triggered by increased growth rates, but not increased size. Two plausible mechanisms for such deceleration can be proposed. One is through mechanical forces and the other is through the action of growth inhibitors; both require to be discussed here. However, a key feature that must first be taken into account is the timing of the deceleration. The fits of the RFI model to the single larva trajectories show that growth deceleration begins shortly after the start. A corresponding conclusion can be made about wing discs based on the pattern of FA during early larval life. What we observe is an increase in FA and subsequent decrease. The increase in FA from zero hours to 15-20 hours must be due to differences in the start time or rate of acceleration between the left and right disc. The subsequent decrease can only be due to regulated deceleration that is triggered at this time. There is scant evidence for candidate molecules or mechanisms that could drive deceleration at this early stage.

Recent work has revealed mechanisms by which growth rates are modulated toward the end of the growth period. A mild increase in hypoxia correlating with disc size was recently proposed as a mechanism for growth deceleration (Zhao et al., 2025). This hypoxia keeps TOR activity in check suggesting a means by which TOR induced biosynthesis doesn’t exceed the demands on available oxygen. Other mechanisms include the effect of ecdysone signalling on the range of morphogens such as Wg and Dpp ((Parker and Struhl, 2020). The decreasing steepness of the Ds gradient is suggested to scale with decreasing cell division rates (Liu et al., 2024). The process of deceleration presented in our study must act alongside these mechanisms. Its primary role is in triggering sustained deceleration in response to the initial acceleration, and so decides rates of deceleration throughout the growth trajectory. A prediction that our study makes is that when discs start growth at a larger initial size (because of overactivation of Dp110 in the embryo), hypoxia would develop at an earlier stage because the size at which hypoxia develops will be reached earlier. Similarly, the establishment of the gradients of the morphogens Dpp and Wg would occur at earlier stages. But once these size correlates are triggered, our study predicts that growth rates (on average across discs), will be determined by history and hence by the time that has passed since the start of growth. It is left to be seen if any of these size correlates override history and initiate a steeper deceleration than that occurs in the controls. Such a study would reveal the effectiveness of each deceleration mechanism, or the ability of each to determine disc growth rates.

Can mechanical forces be responsible for the early deceleration proposed here? The mechanical properties of a tissue dictate their shape and have been shown to determine shape in third instar wing discs (Tripathi et al., 2026). Mechanical properties are likely to correlate with the size of the disc and have been shown to increase within the disc during the third instar stage ((Nienhaus et al., 2009)). In late stage discs, mechanical forces have been shown to alter the growth of clones of tissue ((Shraiman, 2005); (Pan et al., 2016)). In these studies, local increase in compression restricts growth, while stretching boosts growth. During normal growth it is conceivable that such mechanical forces could keep local disturbances in growth rate in check. There have been models proposed that utilize mechanical forces to regulate overall growth rates in an organ-size dependent manner (Aegerter-Wilmsen et al., 2012); (Hecht et al., 2022). But these models have not been experimentally validated. Our model demands that uniform faster rates trigger mechanical feedback keeping the growth rate of the entire disc in check, but in a size independent manner.

It is more conceivable to think of growth inhibitors being made in a growth rate dependent fashion. These could be secreted or could act within cells. If they are secreted, they could play a role in bringing about uniform rates of growth. Whatever the mechanism for growth-rate dependent feedback, our study reveals that to understand the mechanism we should study growth regulation during the first instar of larval life.

### Is size the target of the size determining mechanism?

We show here that extending the larval growth period by using day-night gating, does not stop disc growth. Similar observations have been made by other researchers. Decreasing Ras or Pi3Kinase activity in the prothoracic gland, alters the release of ecdysone, which in turn lengthens the larval period and results in larger flies ((Caldwell et al., 2005). Martin and Morata made mosaic wing discs, in which one compartment consisted of slow growing cells bearing a mutation in a ribosomal protein (the genotype is termed *Minute*) and the other compartment was wildtype ((Morata, 2006). The larval period was extended in these larvae. Interestingly, although the wildtype compartment approached its normal final size earlier, both compartments showed similar frequencies of dividing cells at the end of larval life, just before pupation. These results indicate that cell division continues albeit at a very slow rate, even after normal size is achieved. We cannot rule out the possibility that extending the larval period further would finally stall growth and/or cell division in both compartments. In a study by Parker and Struhl (2020), decreased ecdysone levels leads to slower disc growth rate, but an extended growth period and stalling in disc growth while larvae continue to feed and grow, as ecdysone levels are too low to initiate pupariation.

The question that is being raised in this study, is whether organs need to sense size to stall growth. A true test of whether size is sensed, is when size is altered before the start of the growth period. In this study we show that discs that start large end up making bigger wings, without an alteration in the length of the growth period. If growth stalled at a particular size, then the discs that started at a larger size would have stalled earlier and would not have produced larger wings. Put together, the model proposed in this paper is that deceleration rates are regulated for each organ, such that they occur in proportion to the initial acceleration rate in such a manner that they allow each organ or organism to reach the required size within a given developmental time. With such a model there is no need for size to be sensed.

From an evolutionary point of view, growth rate dependent feedback has the advantage of supporting the principle of parsimony. It suggests that animals that undergo determinate and animals that undergo indeterminate growth achieve their developmental targets through similar mechanisms. This target is to regulate deceleration such that an appropriate size is achieved within the given developmental time without sensing size. What differs in the two kinds of animals is the coordination of growth with other aspects of their life histories. In animals that undergo determinate growth, resources may after the growth period be allocated away from growth and towards survival and reproduction. While in animals that undergo indeterminate growth, resources are utilised towards continued growth albeit at a slow rate.

### Experimental Procedures

#### Fly husbandry and fly stocks

Strains were obtained either from Bloomington Drosophila stock centre, Indiana or DGRC, Vienna. The esg-Gal4-GFP strain was generated by recombination between a fly strain containing a UAS-mCD8-GFP on the second chromosome (BDSC 5137) and a strain bearing P{w+mW.hs=GawB}NP5130 P{UAS-GFP.S65T}T2 P{UAS-GFP::lacZ.nls}2-1 on the second chromosome (DGRC 109-126). Recombinants were selected that expressed GFP on the plasma membrane and not in the nucleus. In the early growth analysis, these recombinants were crossed to yw; UAS-mCD8GFP virgins. In the temperature shift experiments, this recombinant chromosome was combined with tubGal80^ts^ on the third chromosome (BDSC 7017). The yw; UAS-Dp110-CAAX and the yw; UAS-Dp110-KD strains are BDSC 25908 and BDSC 25918, respectively. Where necessary, hemizygous *y* male larvae were distinguished from female first and second instar larvae, with the parents being virgins homozygous for the X-linked yellow mutation *y,* crossed to males that were *y^+^*. The CS strain was cultured in standard fly food containing yeast extract, malt, maize flour and sugar for the wing disc size estimations. All other experiments were performed with flies cultured in food containing 100g dry yeast (Gloripan), 50g Sugar, 50g maize flour, 8g agar, 3 ml Propionic acid and 30ml Nipagin in 1 litre water (based on (Bass et al., 2007). Single larvae were cultured in the latter fly food containing 0.6% Agar instead of 0.8% in shallow capsules placed inside 30mm plastic petri dishes lined with moist tissue. To enable successful pupation, the pupae were transferred to paper cups fixed onto Nipagin soaked paper strips, placed inside long narrow glass vials containing 0.8% Agar and 3% Nipagin.

### Histochemistry

For volume estimation, larvae were dissected in cold PBS and fixed in 4% Formaldehyde, (Electron microscopy grade,) in PLP buffer (0.0316M NaH2PO4, 0.0684M NaHPO4, 1.8% Lysine, 0.00025% NaIO4) for 40 mins, and stained in Rhodamine or AlexaFluor488 conjugated Phalloidin (Molecular Probes1:500) using standard protocols. Confocal images were collected using a Zeiss LSM 510 confocal microscope or a Zeiss Axioobserver7 inverted fluorescence microscope fitted with an Apotome. Volume was estimated using AMIRA image analysis software (Visualization Sciences Group) or 3-D slicer.

### Statistical Analysis

Graphs were plotted and statistical tests were performed using Matlab and R software.

## Supporting information

https://we.tl/t-qD1sAhvq2RSW3g1t

