## Supplementary material for "Sensing rate, not size: growth rate-dependent negative feedback achieves the required organ size": https://we.tl/t-qD1sAhvq2RSW3g1t

### Supplemental Information

#### Figure Legends

Supplemental Figure S1: Analysis of variance in wing imaginal disc size during development.

Supplemental Figure S2: Alignment of Canton S wing disc data with that of *escargot-Gal4, UASmCD8-GFP.*

Supplemental Figure S3: Survival of larvae that were imaged for wing disc FA during early development.

Supplemental Figure S4: Simulations to test convergence of volume when $R_0$ is varied

Supplemental Figure S5: Simulations to test convergence of volume, when $A_0$ is varied

Supplemental Figure S6: Simulations to test convergence of volume, when $I_0$ is varied

Supplemental Figure S7: Growth trajectories of isogenized larvae cultured individually at 29°C.

#### Supplementary Information, Tables and Statistical analysis

Supplemental Information Sheet S1: Analysis of wing disc sizes during the larval period

Supplemental Information Sheet S2: Analysis of Heteroscedasticity, 25°C Canton S wing disc size data.

Supplemental Information Sheet S3: Analysis of Heteroscedasticity, 29°C Canton S wing disc size data.

Supplemental Information Sheet S4: Analysis of Heteroscedasticity, 25°C Canton S combined with esg-Gal4-gfp wing disc size data.

Supplemental Information Table S1: Lag in the start of growth of the wing imaginal discs at 25°C.

Supplemental Information Table S2: Parameter values obtained by fitting mathematical models to 25°C and 29°C Canton S wing disc size data

Supplemental Information Table S3: Fitting RFI model to female larvae data.

### Supplemental Figure Legends

**Supplemental Figure S1: Analysis of variance in wing imaginal disc size during development.** Wing discs were dissected from larvae at different developmental time points. The graphs here represent an analysis of variance in this data. Graphs A-D and E-L represent data from larvae cultured at 29°C and 25°C respectively. The x-axes of A-D, E-H and I-L are aligned with the x-axis of D, H and I respectively and show hours after larval hatching from the embryo, (hAH). Graphs A-H represent data obtained from Canton-S larvae cultured in MYC food (see materials and methods). Graphs I-L represent a composite of data shown in E-H and data obtained from escargot-Gal4, UAS-mCD8GFP larvae grown in Yeast rich food (see materials and methods). Graphs A, E and I are semi-log scatter plots of wing disc volume (in $\mu m^3$, Y-axes are aligned with that of A); B, F and J show fits (black lines) of quadratic polynomials to log(volume) vs hAH (Y-axes aligned with that of B); the squares of the residuals from these fits are shown in C, G and K (Y axes aligned with that of C). Data points in A, E and I are color coded according to experiment date (see text); each point in the above-mentioned graphs represents one wing disc (n=1021 in A-C), 1134 in E-G and 1498 in I-L). Graphs D, H and L represent the analysis of residual variance (see text). The residuals shown in graphs C, G and K were binned into 10-hour bins and the variance calculated for each bin is shown as filled circles in D, H and L respectively. The size of the circles is proportional to the number of data points (wing discs). The curves depict the decrease in variance ($f(x)=e^{2cx}$), with 95% confidence intervals shown as dotted lines and the coefficients of variance c shown in the legend. The analysis of fixed (red in D and blue in H and L) and random effect (yellow in D and magenta in H and L) models are superimposed in the same graphs for easy comparison.

**Supplemental Figure S2: Alignment of Canton S wing disc data with that of *esgargot-Ga4, UAS-mCD8GFP.*** To estimate wing disc sizes during the first quarter of larval life, Z-stack images were taken from fixed larvae expressing membrane targeted GFP (mCD8GFP) under the indirect control of the *escargot* promoter (referred to as esg-Ga4-mCD8GFP). Virgin females carrying the *yellow (y)* mutation were crossed to males that were $y^+$. This scheme enabled the selection of y+/- female larvae, bearing dark mouth hooks. Hemizygous y- males bear pale brown mouth hooks. Only female larvae were dissected and fixed to expose the wing discs. Scatter plot on the left (**A)** displays log(volume) versus developmental time in hours after hatching (hAH). The sizes of the esg-Ga4-mCD8GFP wing discs (green dots) from later stages aligns with the size of wing discs from the Canton S strain (blue dots).

The black curve represents a quadratic polynomial fit to the combined data, the same fit as shown in Figure 1J. Since these experiments were carried out in a different laboratory, from that where the CantonS wing discs were prepared, the food used differed. Thus, the contribution of the change in food and the change in genotype needed to be examined. The deviation of the data from the fit is shown here in **B** as squared residuals. The analysis of variance shown here in **C,** shows that when food/genotype is regarded as a random effect, the high variance at the start of the growth period (heteroscedasticity) is reduced but is not removed.

**Supplemental Figure S3: Survival of larvae that were imaged for wing disc FA during early development.** A scatter plot of the FA index of wing discs for each larva is shown in **A** against time of development as hours after hatching from the embryo, hAH. Esg-Gal4-GFP larvae were immobilised using Desflurane and Apotome images were taken of the left and right wing discs. Each larva was then put back into soft agar food inside single larva culture capsules and allowed to develop until adulthood. A proportion of the larvae did not survive due to handling. The box plot in **B** shows that the survival was not affected by FA index. The mean FA index of survivors and dead larvae was 0.122 and 0.118 respectively with a p value for a Welch two sample test of 0.85.

**Supplemental Figure S4: Simulations to test convergence of volume when $R_0$ is varied**

Random variation was introduced in the initial values of R. The names of the mathematical models are as depicted above each column (see text for expansion of the names). The graphs **A** – **H**, shown here are the same as those shown in Figure 2**A-H.** Two additional panels are added to that the RFA model brings about variation in k1*A (**K**) and the RFI model brings about variation in k2*I (**P**). The Y axes scales are the same as that used in Figure 2, for ease of comparison. Where necessary, details are shown in the insets. Since the k2*I values rise higher than these axes in M, N and O, insets are used to show the entire graphs. The k1*A values rise higher than the scale in **K** and therefore an inset is used to show the entire graph.

**Supplemental Figure S5: Simulations to test convergence of volume when $A_0$ is varied**

Random variation was introduced in the initial values of A. The names of the mathematical models are as depicted above each column (see text for expansion of the names). The graphs **A-L** shown here are the same as graphs shown in Figure 2I-T. The additional panel **M-P** shown here reveals the changes in the k2*I curves that are brought about through the CT and RFI models, when variation in $A_0$ is introduced. The Y axes scales are the same as that used in Figure 2, for ease of comparison. Where necessary, details are shown in the insets. Since the k2*I values rise higher than these scales in M, N and O, insets are used to show the entire graphs. The k1*A values rise higher than the scale in **K** and therefore an inset is used to show the entire graph.

**Supplemental Figure S6: Simulations to test convergence of volume when $I_0$ is varied**

Random variation was introduced in the initial values of I. The names of the mathematical models are as depicted above each column (see text for expansion of the names). The initial values, representing the start of the growth period were obtained from the backward simulations depicted in Figure 2 (dashed lines in Figure 2, see text). The curves representing the highest initial value of $R_0$ are depicted as a black curve. The x-axes for each column align with that of the x-axis of the graph in the bottommost row (Q-T). The Y-axes for graphs in each row align with the Y-axis of the graph on the left. Variation in $I_0$ induces variation in the R curves (**E-H**) and the V curves (**A-D**) for all models. Variation in k1*A is induced with the RFA model. The variation in the initial values was introduced using a random number generator, ensuring that the coefficient of variance (cv) was comparable for all models ($I_0$ at approximately 0.28). The Y axes scales are the same as that used in Figure 2, for ease of comparison. Where necessary, details are shown in the insets. Since the k2*I values rise higher than these scales in M, N and O, insets are used to show the entire graphs. The k1*A values rise higher than the scale in **K** and therefore an inset is used to show the entire graph. Note that the R curves do not cross in the middle of the simulation period in any of the models and correspondingly, the volume curves do not show convergence at the end of the simulation period.

**Supplemental Figure S7: Growth trajectories of isogenized larvae cultured individually at 29°C.** Isogenized larvae were cultured in single larvae culture vials at 29°C, they were immobilized every four hours and volume was estimated from the bright field images taken (see materials and methods), shown here in A as a semi-logarithmic plot of volume versus hours after hatching (hAH). The RFI mathematical model was fit to this data and is shown in graphs B (volume), C (R) and E (I). The graphs are color coded according to the starting value of R (Cyan, red, blue, magenta and green; highest to lowest initial R values). D represents the derivatives of the curves shown in C. The curves reach their minima and then start to increase again. The minimum of each curve is referred to as the peak deceleration value (PDV, arrow in D). Charts F to H represent regression analysis between starting growth rate, starting volume and PDV, with the value of $R^2$, the Regression coefficient indicated. The x-axes of graphs A-D align with that of D.

**Supplementary Information Sheet S1: Analysis of wing disc sizes during the larval period.**

We collected 0-1hr old synchronized egg lays from large population cages containing 1-2 day old Canton S flies. The larvae were collected upon hatching and seeded into culture vials at fixed densities and dissected at regular intervals from the beginning of the second instar until the wandering stage, which marks the end of the larval stage. The eggs and larvae were maintained at either 25°C or 29°C. The discs were stained with Rhodamine conjugated to Phalloidin and the volume was estimated from confocal image stacks. From the late second instar stage onwards, the gonads are visible in male larvae and hence only female larvae were dissected from this stage onwards. Unfortunately, this precaution could not be taken for the early second instar larvae. To address this issue and also examine disc size during the first instar we dissected larvae expressing membrane targeted GFP in imaginal tissue, henceforth referred to as esg-Gal4-GFP larvae (see below). This entire exercise (from egg-lay to larval dissections, regarded as one experiment) was repeated multiple times, to obtain sufficient representation of all stages of growth. Time of larval wandering and pupation was monitored and found to be comparable between experiments. The results are presented in Supplemental Figure S1A (29°C data) Supplemental Figure S1E (25°C CS data) and Supplemental Figure S 1I (25 CS and esg-Gal4-GFP data) as scatter plots of volume versus developmental time (time after hatching from the egg, AH). Each experiment is represented by a different colour. The sizes of discs dissected from larvae from each individual culture vial fall in vertical clusters (for example, see circled clusters in Supplemental Figure S1A, E). It can be observed that until approximately 70 hours after hatching (AH), the vertical clusters from adjacent time values do not align. The size of the smallest disc in a vertical cluster is very often larger than the largest disc from a neighbouring cluster. These volume discrepancies decrease towards the end of the observation period.

The imaginal discs of first instar larvae are too small to be dissected out of the larva. But the discs are easily visible in esg-Gal4-GFP larvae using a fluorescence microscope. Esg-Gal4-GFP larvae which express mCD8GFP under the indirect control of the *escargot* promoter were used (Hayashi et al., 1993); (Goto and Hayashi, 1999). Virgin females of the genotype *yw; UAS-mCD8GFP* were crossed to *w; esg-Gal4, UASmCD8GFP* males. In the resulting progeny larvae, males display brown mouth hooks, because they are hemizygous for the *y* mutation and can be distinguished from *y*/+ heterozygous female larvae. The anterior parts of the female larvae were first fixed in 4% Paraformaldehyde in PBS and then slit open on the ventral sides and mounted in a glycerol based mountant containing antifade. For these experiments the larvae were cultured at 25°C. As described in materials and methods of the main section, the food was changed for these experiments.

To understand the trend in the overall trajectory, quadratic polynomial equations were fit to the data (see Figure S1B, F and J). There was an insignificant difference in the correlation coefficient, $R^2$ and residuals between a quadratic and a cubic polynomial fit to the data sets at both temperatures (summarized in Supplemental information sheets S2, S3 and S4). Therefore, quadratic curve fits were chosen for further analysis, as they represent the simpler model. These fits represent the average disc size at any given developmental time. The squares of the residuals ($(Y\text{-fit})^2$, Figure S1C, G and K) provide a visual representation of the deviation of individual disc sizes from the polynomial fit and confirm that the deviation of disc volume is higher earlier in development and decreases towards the end of this developmental period. To test the statistical significance of this trend we used a generalized least squares method to fit constant variance (homoscedasticity) and non-constant variance (heteroscedasticity) models to the error structure of our data (J. Pinheiro and Bates, 2000). Two models of heteroscedasticity were tested, a power function ($f(x)=x^{2c}$, where c is the variance function coefficient) and an exponential function ($f(x)=b^2e^{2cx}$, where c is the variance function coefficient). A likelihood ratio test and Akaike information criterion showed that both the heteroscedasticity models fit better than the homoscedasticity model for both temperature data sets ($p<0.0001$, summarized in Supplemental Information sheets S2, S3 and S4). Thus, it is quite clear that the variance of the residuals about an average fit or trajectory is not constant. The exponential variance model is more appropriate than the power variance model (see AIC units in Supplemental Information sheets S2, S3 and S4). The variance function coefficient is negative at both temperatures, indicating a decrease in variance (see blue and red line graphs in Figure S1D, H and L and the indicated coefficients, c).

Thus, it is clear that there is greater deviation from an average trajectory or size trend earlier in development. Some of this variance could have arisen because the data was pooled from multiple experiments. To account for this possibility, a linear mixed effects model (J. C. Pinheiro and Bates, 2000) was fit to the residuals, whereby the date of the start of the experiment was taken as a random effect and the polynomial fit as the fixed effect. The date of the start of the experiment serves as a proxy for differences in culture conditions between experiments. The reduction of variance is less steep at both temperatures in the mixed effect than in the fixed effect variance models (see orange and magenta curves in Figure S1D, H and L). AIC units are also much reduced in the mixed effects model (see Supplemental Information sheet S2, S3 and S4). Thus, differences between individual experiments contribute a significant random effect in both the temperature data sets. Note that squared residuals show more of a spread in early development when the CS data is combined with the esg-Gal4-GFP data (see blue dots between 0-30h AH in Figure S1K). The residual variance at the start is much higher in the combined data set than in the CS data set (compare coefficients, c between Figure S1L and 1H). This is particularly notable as only female esg-Gal4-GFP larvae were analysed.

As revealed in materials and methods, the esg-Gal4-GFP larvae were grown using a different food recipe from that used for the CS larvae. Despite this change in food and genotype of the larvae, we found that the volume of later time points aligned well with the data from CS larvae, (Supplemental Figures S1I and S2) and the variance at the start remains high in a mixed effect model, in which food was used as a random effect (see Supplemental Figure S2C).

The observation that the heteroscedasticity remains significant in the mixed effects models is an indication that it is due to more than variation in conditions between different experiments. One possibility could be that there are minor culture differences between vials within each experiment. Larvae within a culture vial could influence the rate of growth of other larvae in the same vial, possibly by influencing the overall level of moisture in the vial. These collective growth rates could differ between one vial and another. Another source of such variation could be developmental stochasticity. Developmental stochasticity in the start of growth is manifest as fluctuating asymmetry (FA) in the sizes of left and right wings. This analysis is represented in the main text and the details are represented below.

**Image Acquisition and analysis of FA in wing imaginal discs:**

The esg-Gal4-GFP larvae were used for this analysis. Discs were imaged from live larvae using an Zeiss inverted fluorescence Axio-Observer 7 microscope, fitted with Apotome. To enable image acquisition, the esg-Gal4-GFP larvae were immobilized using Desflurane, using a device called Flexiheight (described elsewhere). A small amount of pressure was applied on the larva to bring the disc closer to the microscope lens. Larvae survive such treatment when only one side is imaged and the likelihood of survival depends on how much pressure is applied and how much Desflurane is delivered. Image acquisition of discs on both sides requires repeated anaesthetization over a short interval of less than 11 minutes reducing their chances of survival. Hence, each larva was imaged at only one time point and then placed back inside the food. To obtain sufficient numbers, disc pairs were considered in the analysis, even when the larva did not survive to adulthood. The scatter plot in Supplemental Figure S3 shows that even larvae that show higher disc FA were able to survive to adulthood, and the FA of survivors is not significantly different from the FA of those that died (mean FA being 0.122 and 0.118, respectively with p value from a Welch two sample test of 0.85), suggesting that death is due to handling rather than high FA. Since the volume of discs at these early stages is very small, slight errors in measurement could obscure the analysis of FA. Thus, each image was analysed at least twice and measurement error was calculated as the difference between two measurements divided by the mean measurement value. The analysis was done blind, with the discs numbered randomly removing any indication of stage or larval identity. In the first two sets of images, 59 out of 138 discs revealed a measurement error of less than 0.05. The remaining discs were analysed again. From these three sets of measurements, measurement pairs for 136 discs were obtained with a measurement error of less than 0.06. These values were considered in the analysis.

**Supplemental Information Sheet S2: Analysis of Heteroscedasticity, 25°C Canton S wing disc size data.** The statistical analysis of variance of the 25°C volume data versus developmental time (hours) shows that deviation from the average trajectory decreases significantly with developmental time. The analysis was performed using R software (R Core Team, 2026), according to the guidelines of Pinheiro and Bates, 2000.

---

Fit of quadratic polynomial
Call: lm(formula = log(volume) ~ hours + I(hours^2))
Coefficients:

| | Estimate | Std. Error | t value | Pr(>\|t\|) |
|---|---|---|---|---|
| Intercept | 7.36 | 0.07213 | 102.04 | ***$2x10^{-16}$ |
| hours | 0.1235 | 0.002396 | 51.53 | ***$2x10^{-16}$ |
| $hours^2$ | -0.0004528 | 0.00001868 | -24.24 | ***$2x10^{-16}$ |

Residual standard error: 0.306 on 1131 degrees of freedom

Multiple R-squared: 0.954, Adjusted R-squared: 0.9539

F-statistic: 1.173e+04 on 2 and 1131 DF, p-value: < 2.2e-16

---

Fit of cubic polynomial
Call: lm(formula = log(volume) ~ hours + I(hours^2)) + I(hours^3)
Coefficients:

| | Estimate | Std. Error | t value | Pr(>\|t\|) |
|---|---|---|---|---|
| Intercept | 7.823 | 0.1807 | 43.299 | ***$2x10^{-16}$ |
| hours | 0.09658 | 0.009935 | 9.71 | ***$2x10^{-16}$ |
| $hours^2$ | 0.0000151 | 0.0001688 | 0.089 | 0.92874 |
| $hours^3$ | -0.000002487 | $8.9x10^{-7}$ | -2.789 | ** 0.00537 |

Residual standard error: 0.3051 on 1130 degrees of freedom

Multiple R-squared: 0.9543, Adjusted R-squared: 0.9542

F-statistic: 7869 on 3 and 1130 DF, p-value: < 2.2e-16

---

**Since the $R^2$ values and residual standard errors were not different between the quadratic and cubic fits, the simpler quadratic polynomial model was chosen.**

Using the quadratic polynomial fit, homoscedasticity versus heteroscedasticity was tested:

Results of ANOVA analysis of residual variance:

| | Df | AIC units | Anova test | Likelihood ratio | p-value |
|---|---|---|---|---|---|
| Constant variance | 4 | 578.77 | | | |
| Variance changing as a power function | 5 | 563.62 | Constant vs power | 17.16 | <0.0001 |
| Variance changing as an exponential function | 5 | 550.24 | Constant vs exponential | 30.53 | <0.0001 |

AIC units were lowest for the exponential variance model (less by more than 10 AIC units). Therefore, the exponential variance model was preferred. Exponential coefficient for heteroscedasticity: -0.00652. The coefficient is negative. Therefore, the residual variance decreases with increasing values of the x-axis (which denotes developmental time).

---

Introduction of date of experiment as a random effect:

Results of ANOVA analysis:

| | Df | AIC units | ANOVA test | |
|---|---|---|---|---|
| | | | Likelihood ratio | p-value |
| Variance changing as an exponential function, no random effect | 5 | 550.24 | | |
| Variance changing as an exponential function, date as random effect | 6 | 354.50 | 197.74 | <0.0001 |

The AIC units are much lower when date is introduced as a random effect. Therefore, date has a significant effect on the residual variance. Exponential coefficient of heteroscedasticity with date as random effect: -0.00648

**Supplemental Information Sheet S3: Analysis of Heteroscedasticity, 29°C Canton S wing disc data.** The statistical analysis of variance of the 29°C volume data versus developmental time (hours) shows that deviation from the average trajectory decreases significantly with developmental time. The analysis was performed using R software (R Core Team, 2026), according to the guidelines of Pinheiro and Bates, 2000.

---

Fit of quadratic polynomial
Call: lm(formula = log(volume) ~ hours + I(hours^2))
Coefficients:

| | Estimate | Std. Error | t value | Pr(>\|t\|) |
|---|---|---|---|---|
| Intercept | 7.127 | 0.08092 | 88.07 | ***$2x10^{-16}$ |
| hours | 0.1546 | 0.003102 | 49.83 | ***$2x10^{-16}$ |
| $hours^2$ | -0.0007158 | 0.00002756 | -25.97 | ***$2x10^{-16}$ |

Residual standard error: 0.3413 on 1018 degrees of freedom

Multiple R-squared: 0.9435, Adjusted R-squared: 0.9434

F-statistic: 8506 on 2 and 1018 DF, p-value: < 2.2e-16

---

Fit of cubic polynomial
Call: lm(formula = log(volume) ~ hours + I(hours^2)) + I(hours^3)
Coefficients:

| | Estimate | Std. Error | t value | Pr(>\|t\|) |
|---|---|---|---|---|
| Intercept | 8.126 | 0.2068 | 39.292 | ***$<2x10^{-16}$ |
| hours | 0.0914 | 0.01245 | 7.341 | ***$4.32x10^{-13}$ |
| $hours^2$ | 0.0004883 | 0.0002315 | 2.109 | *0.0352 |
| $hours^3$ | -0.000007 | $1.3x10^{-6}$ | -5.237 | ***$1.98x10^{-7}$ |

Residual standard error: 0.3369 on 1017 degrees of freedom

Multiple R-squared: 0.945, Adjusted R-squared: 0.9499

F-statistic: 5827 on 3 and 1017 DF, p-value: < 2.2e-16

---

**Since the $R^2$ values and residual standard errors were not different between the quadratic and cubic fits, the simpler quadratic polynomial model was chosen.**

Using the quadratic polynomial fit, homoscedasticity versus heteroscedasticity was tested:

Results of ANOVA analysis of residual variance:

| | Df | AIC units | Anova test | Likelihood ratio | p-value |
|---|---|---|---|---|---|
| Constant variance | 4 | 746.65 | | | |
| Variance changing as a power function | 5 | 733.45 | Constant vs power | 14.83 | ***0.0001 |
| Variance changing as an exponential function | 5 | 719.56 | Constant vs exponential | 29.09 | ***<0.0001 |

AIC units were lowest for the exponential variance model (less by more than 10 AIC units). Therefore, the exponential variance model was preferred. Exponential coefficient for heteroscedasticity: -0.00838. The coefficient is negative. Therefore, the residual variance decreases with increasing values of the x-axis (which denotes developmental time).

---

Introduction of date of experiment as a random effect:

Results of ANOVA analysis:

| | Df | AIC units | ANOVA test | |
|---|---|---|---|---|
| | | | Likelihood ratio | p-value |
| Variance changing as an exponential function, no random effect | 5 | 719.56 | | |
| Variance changing as an exponential function, date as random effect | 6 | 376.20 | 374.92 | ***<0.0001 |

The AIC units are much lower when date is introduced as a random effect. Therefore, date has a significant effect on the residual variance. Exponential coefficient of heteroscedasticity with date as random effect: -0.00496

**Supplemental Information Sheet S4: Analysis of Heteroscedasticity, 25°C Canton S combined with esg-Gal4-gfp wing disc size data**. The Canton S cultures and the esg-Gal4-gfp cultures were reared on MYC and Yeast rich food respectively (see materials and methods, main text). The statistical analysis of variance versus developmental time (hours) shows a similar trend to the Canton C data alone, but with an increased negative coefficient of residual variance.

Fit of quadratic polynomial
Call: lm(formula = log(volume) ~ hours + I(hours^2))
Coefficients:

| | Estimate | Std. Error | t value | Pr(>\|t\|) |
|---|---|---|---|---|
| Intercept | 6.507 | 0.0362 | 179.56 | ***$2x10^{-16}$ |
| hours | 0.1479 | 0.001454 | 101.72 | ***$2x10^{-16}$ |
| $hours^2$ | -0.0006175 | 0.00001326 | -46.58 | ***$2x10^{-16}$ |

Residual standard error: 0.3641 on 1497 degrees of freedom

Multiple R-squared: 0.9716, Adjusted R-squared: 0.9715

F-statistic: 2556 on 2 and 1495 DF, p-value: < 2.2e-16

Fit of cubic polynomial
Call: lm(formula = log(volume) ~ hours + I(hours^2)) + I(hours^3)
Coefficients:

| | Estimate | Std. Error | t value | Pr(>\|t\|) |
|---|---|---|---|---|
| Intercept | 6.489 | 0.05241 | 123.83 | ***$<2x10^{-16}$ |
| hours | 0.1495 | 0.003827 | 39.073 | ***$<2x10^{-16}$ |
| $hours^2$ | -0.000654 | 0.00008017 | -8.157 | ***$7.19x10^{-16}$ |
| $hours^3$ | 0.0000002264 | $4.9x10^{-7}$ | 0.461 | 0.645 |

Residual standard error: 0.3649 on 1494 degrees of freedom

Multiple R-squared: 0.9716, Adjusted R-squared: 0.9715

F-statistic: 1703 on 3 and 1494 DF, p-value: < 2.2e-16

**Since the $R^2$ values and residual standard errors were not different between the quadratic and cubic fits, the simpler quadratic polynomial model was chosen.**

Using the quadratic polynomial fit, homoscedasticity versus heteroscedasticity was tested:

Results of ANOVA analysis of residual variance:

| | Df | AIC units | Anova test | Likelihood ratio | p-value |
|---|---|---|---|---|---|
| Constant variance | 4 | 1277.206 | | | |
| Variance changing as a power function | 5 | 1197.286 | Constant vs power | 81.92 | ***<0.0001 |
| Variance changing as an exponential function | 5 | 1114.81 | Constant vs exponential | 164.40 | ***<0.0001 |

AIC units were lowest for the exponential variance model (less by more than 10 AIC units). Therefore, the exponential variance model was preferred. Exponential coefficient for heteroscedasticity: -0.00985. This value has a larger negative value that that for the Canton S data alone. The likelihood ratios when comparing the heteroscedasticity models with constant variance models are higher than the same with the Canton-S data alone. The coefficient is negative. Therefore, the residual variance decreases with increasing values of the x-axis (which denotes developmental time).

Introduction of date of experiment as a random effect:

Results of ANOVA analysis of exponential function with and without date as random effect:

| | Df | AIC units | ANOVA test | |
|---|---|---|---|---|
| | | | Likelihood ratio | p-value |
| Variance changing as an exponential function, no random effect | 5 | 1114.81 | | |
| Variance changing as an exponential function, date as random effect | 6 | 826.99 | 289.83 | ***<0.0001 |

The AIC units are much lower when date is introduced as a random effect. Therefore, date has a significant effect on the residual variance. Exponential coefficient of heteroscedasticity with date as random effect: -0.0088.

Introduction of food / genotype as a random effect:

Results of ANOVA analysis:

| | Df | AIC units | ANOVA test | |
|---|---|---|---|---|
| | | | Likelihood ratio | p-value |
| Variance changing as an exponential function, no random effect | 5 | 1114.81 | | |
| Variance changing as an exponential function, food/genotype as random effect | 6 | 1063.38 | 53.43 | ***<0.0001 |

The AIC units are much lower when food/genotype is introduced as a random effect. Therefore, food or genotype has a significant effect on the residual variance. However, the AIC units are much higher here than when date is used as random effect. Thus, variation between experiments has a more significant effect on residual variance than genotype or food. Exponential coefficient of heteroscedasticity with food/genotype as random effect: -0.0080

Supplemental Table S1: Lag in the start of growth of the wing imaginal discs at 25°C.

| hours AH | 0 hrs | 3.35 hrs | 4.15 hrs | 6 hrs | 6.15 hrs |
|---|---|---|---|---|---|
| Volume ($\mu m^3$) | 1405 ± 343 SD | 1375 ± 289 SD | 1296 ± 230 SD | 1108 ± 157 SD | 2125 ± 497 SD |
| p value | | 0.83 | 0.40 | 0.016 | 0.000731 |
| Range ($\mu m^3$) | 935 to 2136 | 890 to 1644 | 1032 to 1742 | 882 to 1441 | 1053 to 3016 |

The mean volumes are shown of wing imaginal discs dissected at the given developmental times shown as hours after hatching of the larva from the egg (hoursAH). The discs are from esg-Ga4-GFP larvae and express mCD8GFP in imaginal tissue. The p values are of student t-tests performed to compare the means of each sample set with the mean volume at 0 hours. There is no significant difference in the means of the first three samples. The two samples at 6 hrs and 6.15 hrs significantly differ from the sample at 0 hrs, one of them shows a decrease in mean and the other shows an increase in the mean.

Supplemental Information Table S2: Parameter values obtained by fitting mathematical models to 25°C and 29°C Canton S wing disc size data

| | 25°C | | | | 29°C | | | |
|---|---|---|---|---|---|---|---|---|
| | NCR | CT | RFA | RFI | NCR | CT | RFA | RFI |
| $V_0$ | 19668 | 21287 | 21105 | 20292 | 21866 | 22031 | 21720 | 22273 |
| $R_0$ | 0.092 | 0.0852 | 0.085 | 0.088 | 0.102 | 0.103 | 0.102 | 0.0996 |
| $A_0$ | 0.004 | 0.0037 | 0.005 | 0.005 | 0.005 | 0.005 | 0.015 | 0.007 |
| $I_0$ | 0.005 | 0.0052 | 0.005 | 0.005 | 0.005 | 0.007 | 0.007 | 0.002 |
| k1 | 0.892 | 0.231 | 0.297 | 0.294 | 0.787 | 0.389 | 0.348 | 0.274 |
| k2 | 0.690 | 0.18 | 0.233 | 0.173 | 0.594 | 0.241 | 0.352 | 0.326 |
| k3 | 1.613 | 1.681 | 1.611 | 1.540 | 1.691 | 1.163 | -0.721 | 0.908 |
| k4 | 1.743 | 1.811 | 1.582 | 1.650 | 1.721 | 1.29 | -0.954 | 0.967 |
| k5 | 3.610 | 1.491 | 1.391 | 1.414 | 3.587 | 1.184 | 1.276 | 1.093 |
| k6 | 3.578 | 1.432 | 1.333 | 1.472 | 3.535 | 1.123 | 1.22 | 1.115 |
| k7 | 1.408 | 0.335 | 1.593 | 1.528 | 1.522 | 1.628 | 3.112 | 1.16 |
| k8 | - | 1.559 | - | - | - | 1.292 | - | - |
| $R^2$ | 0.954 | 0.954 | 0.954 | 0.955 | 0.945 | 0.945 | 0.945 | 0.945 |
| $V_0$ | 3215 | 3558 | 3584 | 3289 | 3334 | 3394 | 3560 | 3495 |
| $R_0$ | 0.0642 | 0.0777 | 0.0764 | 0.0782 | 0.0946 | 0.0883 | 0.0849 | 0.0938 |
| $A_0$ | 0.0641 | 0.0620 | 0.0463 | 0.0504 | 0.0093 | 0.0616 | 0.0496 | 0.0213 |
| $I_0$ | 0.0026 | 0.000729 | 0.0013 | 0.0011 | 0.0018 | 0.0013 | 0.0024 | 0.00039 |

Multiple simulations were performed to fit the models NCR, CT, RFA and RFI to the 25°C and 29°C disc volume data. We found it convenient to fix the starting values for the constants k1 to k8 and $A_0$ and $I_0$. The starting values were 0.3, 0.18, 1.55, 1.55, 2, 1.276, 1.5, 1.5, 0.005 and 0.005 for k1 to k8 and $A_0$ and $I_0$, respectively. At least nine simulations were run for each model, for each data set, with different combinations of starting values for $V_0$ and $R_0$. The above table lists the values obtained for the initial conditions $V_0$ to $I_0$ and the constants k1 to k8, with the best R2 values. The last for rows of the above table display the values obtained in backward simulations (dotted lines in Figure 2), equating to 0hAH.

Some simulations gave negative values for some of the constants. These were not considered. However, note that the both the k3 and k4 values for the RFA simulation for the 29 data have negative values. Thus, in effect the equation becomes the following:

$$RFA: Equation\ (3)\ \frac{dA}{dt} = -k_3A - k_7RA - (-k_4A)$$

$$RFA: Equation\ (3)\ \frac{dA}{dt} = k_4A - k_7RA - k_3A$$

Supplemental Information Table S3: Fitting RFI model to female larvae data

| | 25°C | 29°C |
|---|---|---|
| $V_0$ | 12.77 | 12.5 |
| $R_0$ | 0.073 | 0.09 |
| $A_0$ | 0.003 | 0.013 |
| $I_0$ | 0.005 | 0.001 |
| k1 | 0.618 | 0.192 |
| k2 | 0.072 | 0.335 |
| k3 | 1.557 | 1.143 |
| k4 | 2.184 | 2.228 |
| k5 | 1.951 | 4.34 |
| k6 | 1.94 | 4.345 |
| k7 | 0.94 | 1.308 |
| k8 | - | - |
| $R^2$ | 0.98 | 0.988 |

Multiple simulations were performed to fit the RFI model to the 25°C and 29°C larval volume data. We found it convenient to fix the starting values for the constants k1 to k8 and $A_0$ and $I_0$. The starting values were 0.3, 0.18, 1.55, 2.5, 2, 1.276, 1.5, 0.005 and 0.005 for k1 to k7 and $A_0$ and $I_0$, respectively for the 25°C data and 0.3, 0.6, 1.55, 1.8, 3.57, 3.54, 1.5, 0.005 and 0.005 accordingly for the 29°C data. At least nine simulations were run for each model, for each data set, with different combinations of starting values for $V_0$ and $R_0$. The above table lists the values obtained for the initial conditions $V_0$ to $I_0$ and the constants k1 to k8, with the best $R^2$ values.

Table S5: Fitting RFI model to single larvae data.

| Larva Serial No. | 25°C | | | | | 29°C | | | | |
|---|---|---|---|---|---|---|---|---|---|---|
| | $V_0$ | $R_0$ | $A_0$ | $I_0$ | $R^2$ | $V_0$ | $R_0$ | $A_0$ | $I_0$ | $R^2$ |
| 1 | 13.66 | 0.063 | 0.0072 | 0.0052 | 0.995 | 15.78 | 0.093 | 0.0047 | 0.0014 | 0.997 |
| 2 | 10.03 | 0.089 | 0.0066 | 0.0055 | 0.997 | 13.18 | 0.084 | 0.0066 | 0.0008 | 0.998 |
| 3 | 12.89 | 0.064 | 0.0057 | 0.0021 | 0.990 | 15.86 | 0.086 | 0.0076 | 0.0014 | 0.998 |
| 4 | 10.41 | 0.078 | 0.0048 | 0.0043 | 0.997 | 15.31 | 0.086 | 0.0072 | 0.0020 | 0.998 |
| 5 | 12.50 | 0.072 | 0.0075 | 0.0050 | 0.997 | 15.31 | 0.086 | 0.0072 | 0.0020 | 0.998 |
| 6 | 13.92 | 0.067 | 0.0029 | 0.0038 | 0.997 | 13.70 | 0.088 | 0.0148 | 0.0011 | 0.998 |
| 7 | 14.25 | 0.056 | 0.0047 | 0.0005 | 0.995 | 16.98 | 0.082 | 0.0186 | 0.0016 | 0.999 |
| 8 | 11.94 | 0.077 | 0.0041 | 0.0041 | 0.997 | 15.12 | 0.094 | 0.0195 | 0.0013 | 0.998 |
| 9 | 13.90 | 0.063 | 0.0074 | 0.0047 | 0.995 | 16.37 | 0.086 | 0.0181 | 0.0015 | 0.998 |
| 10 | 12.06 | 0.067 | 0.0009 | 0.0030 | 0.997 | 9.94 | 0.088 | 0.0121 | 0.0009 | 0.997 |
| 11 | 12.47 | 0.078 | 0.0079 | 0.0049 | 0.997 | 11.90 | 0.096 | 0.0061 | 0.0012 | 0.997 |
| 12 | 13.84 | 0.069 | 0.0058 | 0.0039 | 0.998 | 9.92 | 0.093 | 0.0111 | 0.0009 | 0.998 |
| 13 | 17.10 | 0.063 | 0.0070 | 0.0037 | 0.997 | 10.57 | 0.096 | 0.0111 | 0.0009 | 0.997 |
| 14 | 11.69 | 0.069 | 0.0049 | 0.0032 | 0.998 | 10.45 | 0.085 | 0.0080 | 0.0009 | 0.997 |
| 15 | 13.06 | 0.071 | 0.0040 | 0.0040 | 0.999 | 13.09 | 0.090 | 0.0074 | 0.0014 | 0.995 |
| 16 | 14.66 | 0.074 | 0.0042 | 0.0051 | 0.998 | 10.65 | 0.096 | 0.0158 | 0.0012 | 0.997 |
| 17 | 12.73 | 0.060 | 0.0097 | 0.0014 | 0.996 | 11.85 | 0.098 | 0.0149 | 0.0011 | 0.997 |
| 18 | 12.61 | 0.075 | 0.0071 | 0.0047 | 0.998 | 9.98 | 0.098 | 0.0070 | 0.0012 | 0.999 |
| 19 | 14.54 | 0.065 | 0.0040 | 0.0070 | 0.996 | 10.69 | 0.086 | 0.0091 | 0.0008 | 0.996 |
| 20 | 11.25 | 0.078 | 0.0054 | 0.0052 | 0.997 | 12.93 | 0.085 | 0.0070 | 0.0010 | 0.997 |
| 21 | 11.82 | 0.080 | 0.0084 | 0.0053 | 0.996 | 11.32 | 0.090 | 0.0088 | 0.0009 | 0.997 |
| 22 | 11.22 | 0.082 | 0.0084 | 0.0050 | 0.998 | 9.84 | 0.101 | 0.0093 | 0.0009 | 0.996 |
| 23 | 12.88 | 0.080 | 0.0060 | 0.0056 | 0.997 | | | | | |
| 24 | 11.39 | 0.076 | 0.0077 | 0.0048 | 0.998 | | | | | |
| 25 | 15.02 | 0.079 | 0.0044 | 0.0063 | 0.997 | | | | | |
| 26 | 11.75 | 0.080 | 0.0071 | 0.0061 | 0.998 | | | | | |
| 27 | 13.26 | 0.082 | 0.0065 | 0.0067 | 0.997 | | | | | |
| 28 | 14.11 | 0.070 | 0.0074 | 0.0051 | 0.997 | | | | | |
| 29 | 12.14 | 0.073 | 0.0073 | 0.0046 | 0.997 | | | | | |
| 30 | 11.06 | 0.063 | 0.0067 | 0.0022 | 0.990 | | | | | |
| 31 | 11.56 | 0.089 | 0.0064 | 0.0070 | 0.993 | | | | | |
| 32 | 11.27 | 0.077 | 0.0076 | 0.0047 | 0.998 | | | | | |
| 33 | 13.29 | 0.070 | 0.0039 | 0.0049 | 0.998 | | | | | |
| 34 | 13.82 | 0.068 | 0.0074 | 0.0051 | 0.994 | | | | | |
| 35 | 14.47 | 0.082 | 0.0002 | 0.0080 | 0.998 | | | | | |
| 36 | 15.40 | 0.078 | 0.0049 | 0.0067 | 0.992 | | | | | |
| 37 | 12.91 | 0.072 | 0.0055 | 0.0053 | 0.998 | | | | | |
| 38 | 11.73 | 0.073 | 0.0044 | 0.0047 | 0.997 | | | | | |

| | $R^2$ values of linear regression analysis |
|---|---|
| $R_0$ vs PDV | 0.9754 (negative slope) |
| $R_0$ vs Actual starting volume | 0.1271 |
| PDV vs actual starting volume | 0.1081 |
| R at peak deceleration vs $R_0$ | 0.01879 |
| Time at peak deceleration vs $R_0$ | 0.7796 (negative slope) |
| Volume at peak deceleration vs $R_0$ | 0.1596 |
| Volume at peak deceleration vs at start | 0.06899 |

**Supplemental Information Table S4: Regression analysis of fits of RFI model to single larval trajectories at 25°C.** $R_0$ is the initial value for R, obtained for each fit to each larval trajectory. PDV, the peak deceleration value represents the lowest value of $d^2logV/dt^2$, for each trajectory, an indicator of the highest level of deceleration for each larva (see main text). The rate (dlogV/dt) represented by R at this PDV does not correlate with either the starting rate or starting volume. Similiarly the volume of the larva at PDV does not correlate with its starting volume or starting growth rate. The strongest correlation is that of $R_0$ vs PDV and weaker but significant correlation is shown between time at which this PDV is reached and $R_0$, as larvae that start at a faster rate, slow down earlier and faster. The rate of deceleration then eases, as decrease in R results in a decrease in inhibitor synthesis.

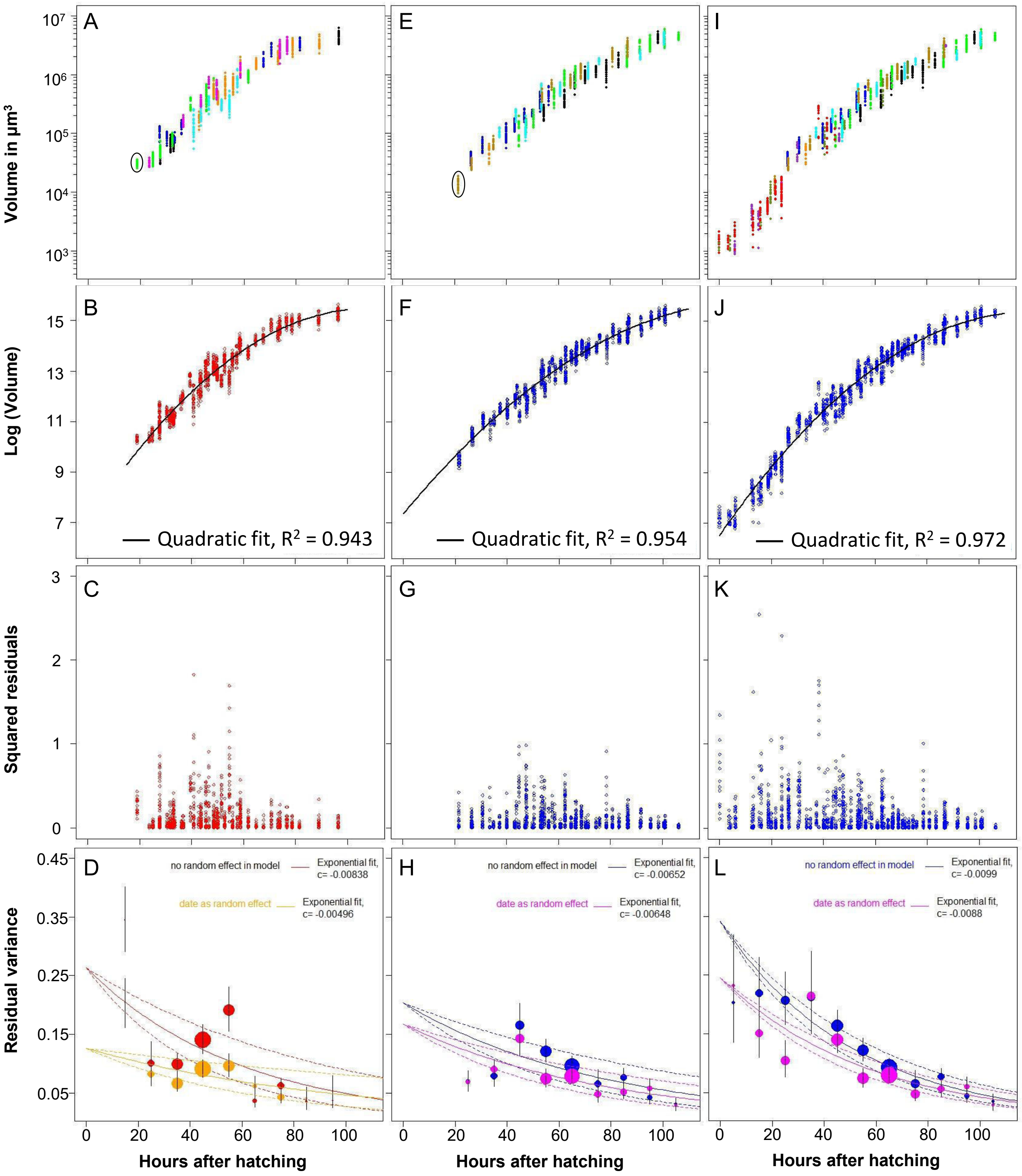


Figure S1: Analysis of variance of wing imaginal size during development

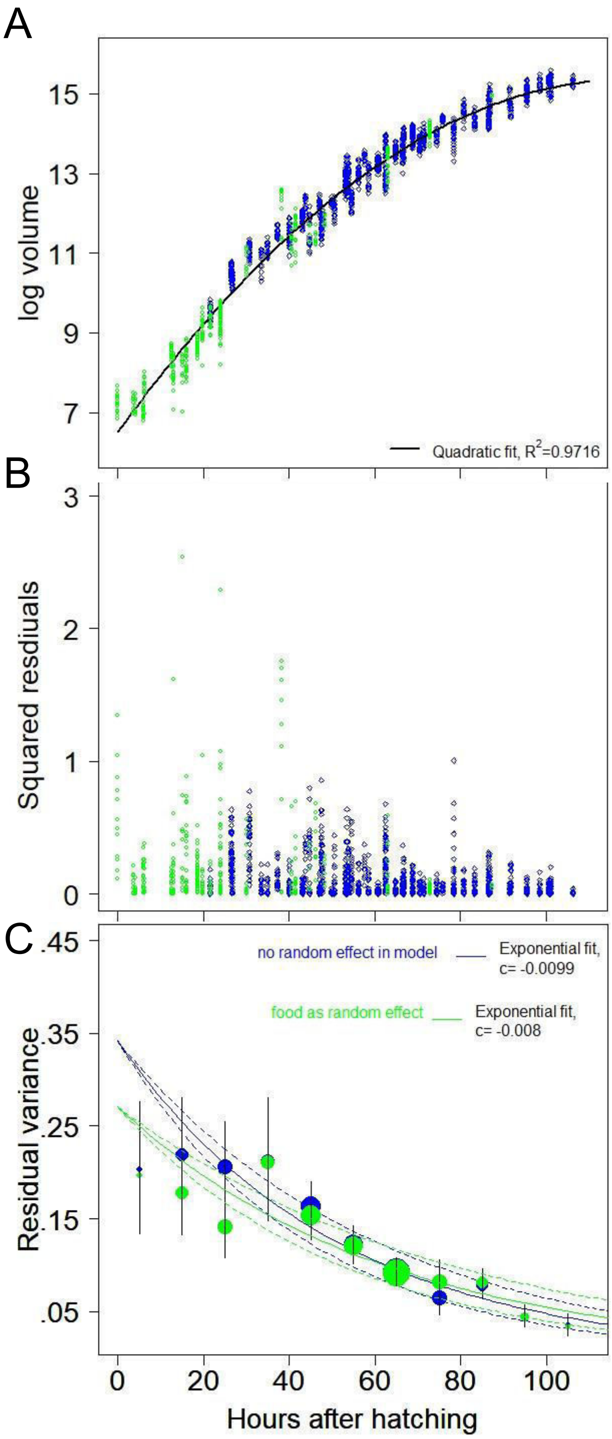


Supplemental Figure S2: Alignement of CantonS wing disc data with that of *escargot-Gal4, UAS-mCD8GFP*

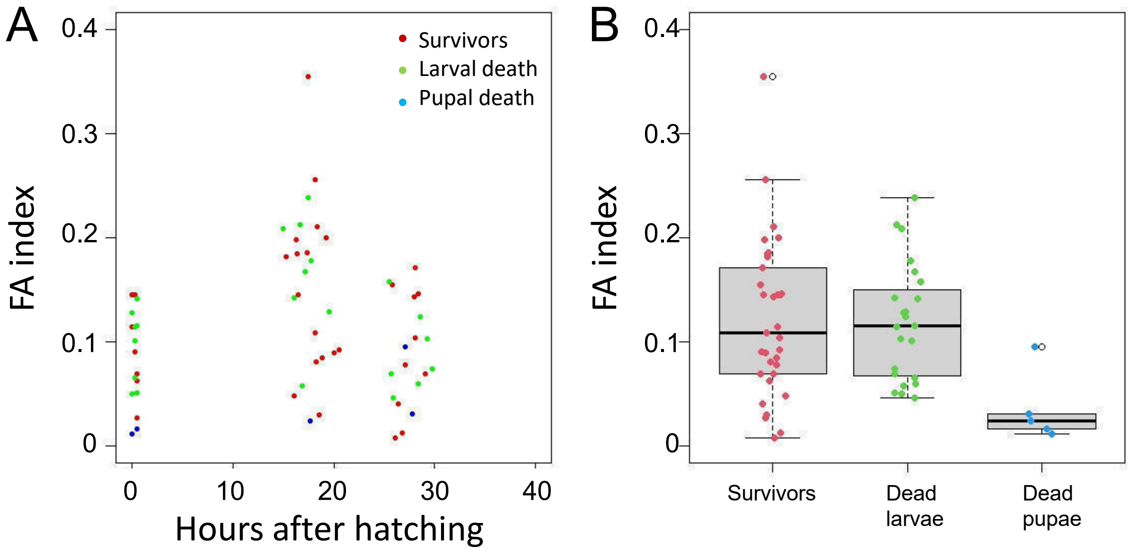


Supplemental Figure S3: Survival of larvae that were imaged for wing disc FA during early development.

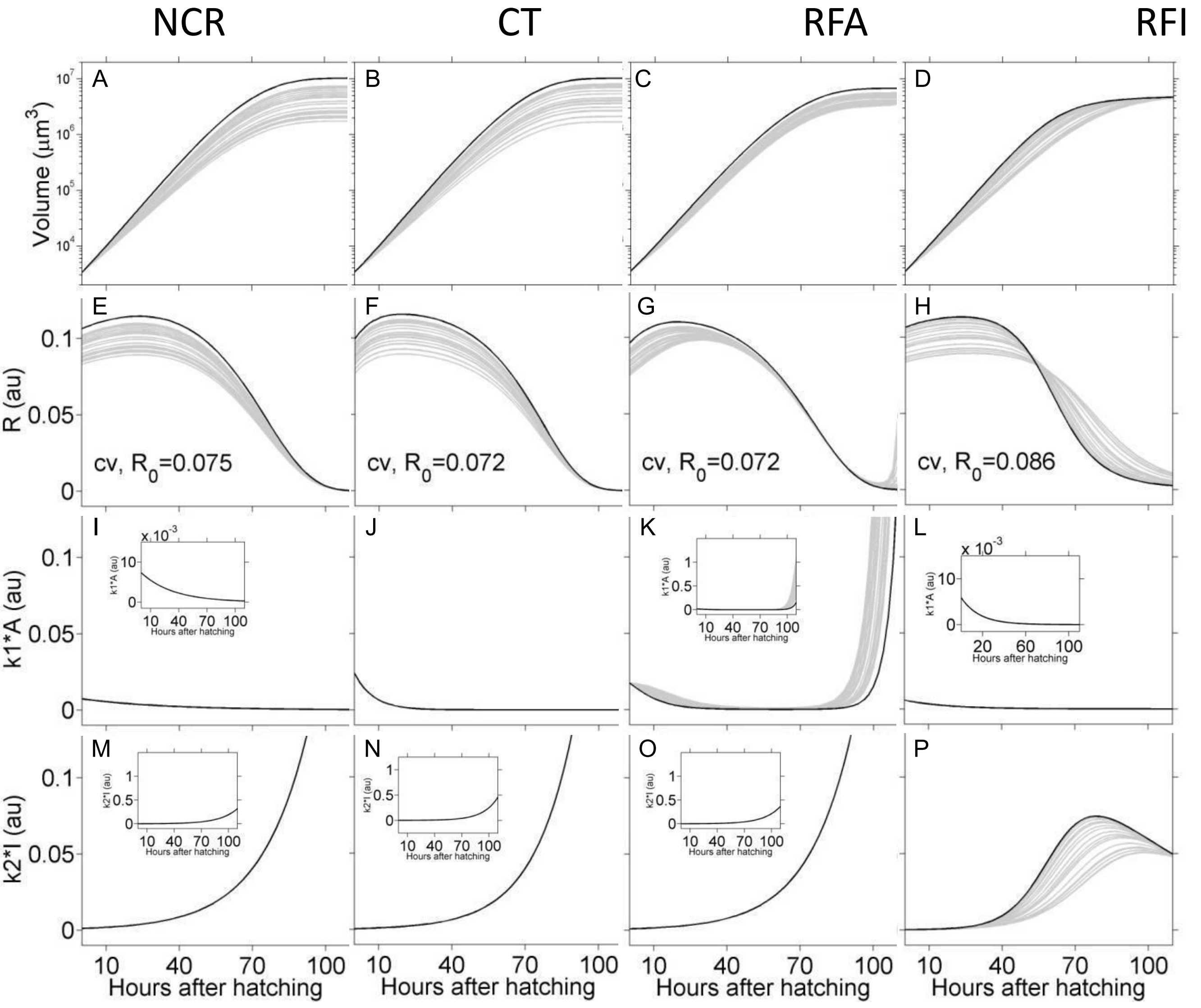


Supplemental Figure S4: Simulations to test convergence of volume when $R_0$ is varied

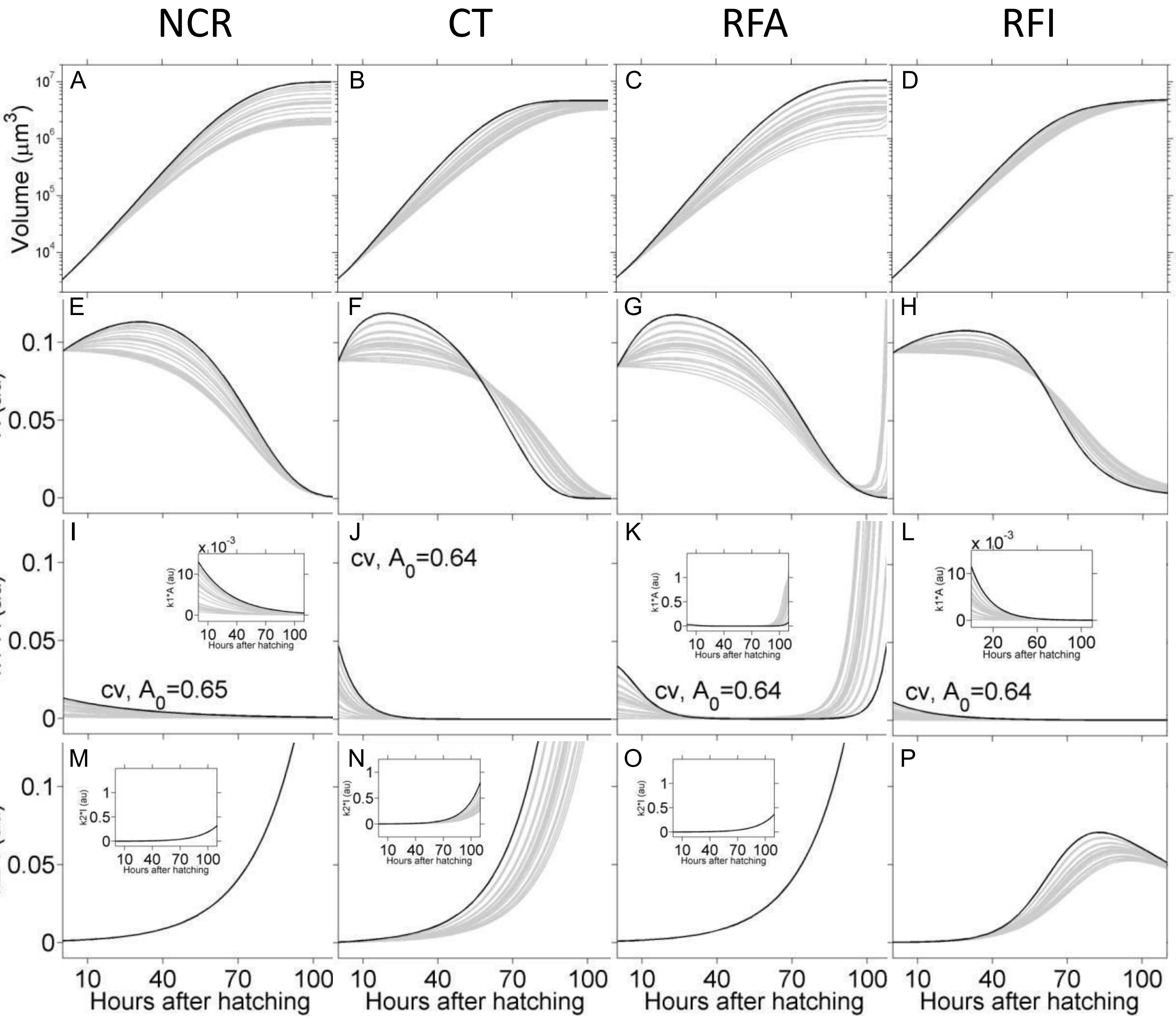


Supplemental Figure S5: Simulations to test convergence of volume, when $A_0$ is varied

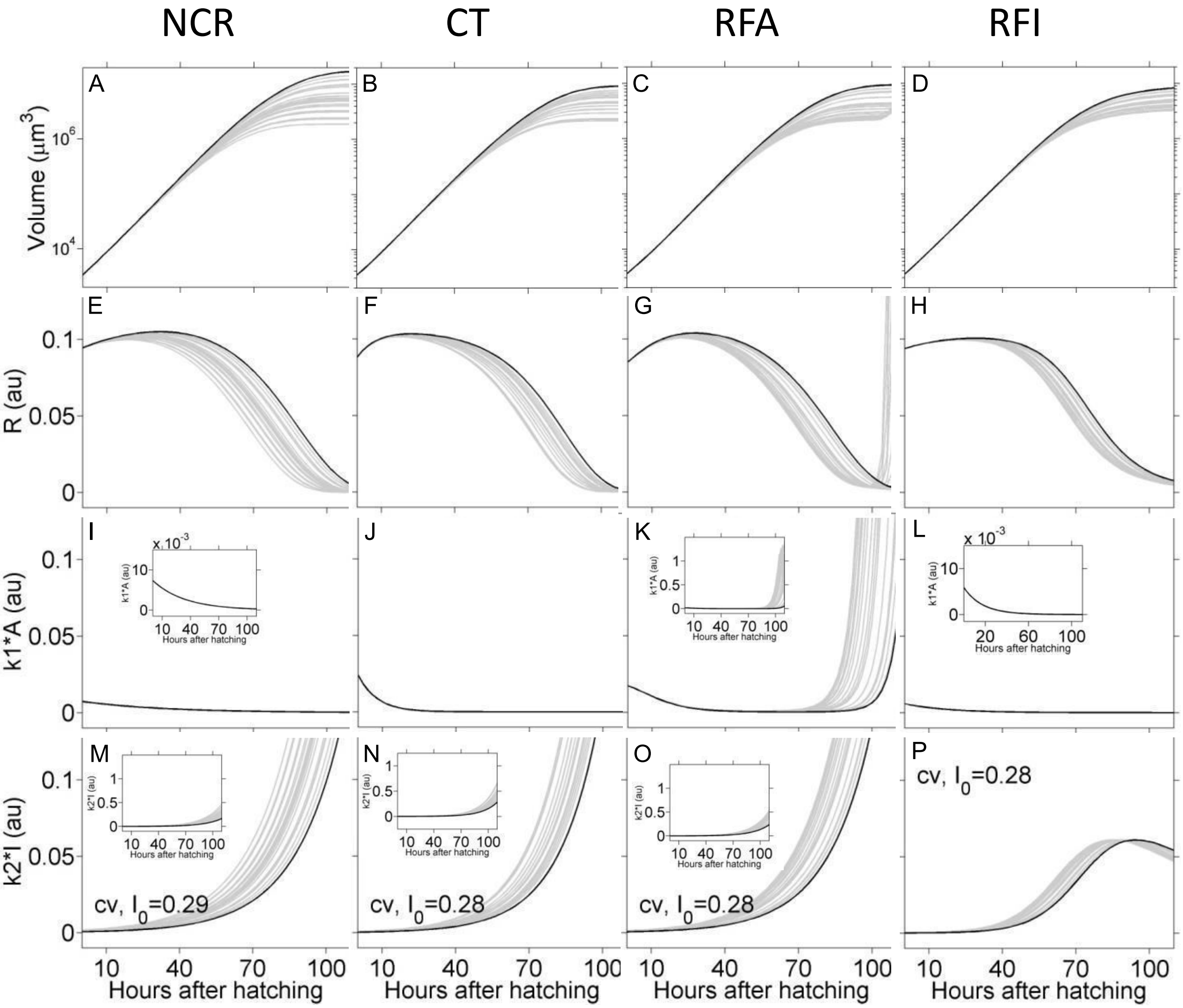


Supplemental Figure S6: Simulations to test convergence of volume, when $I_0$ is varied

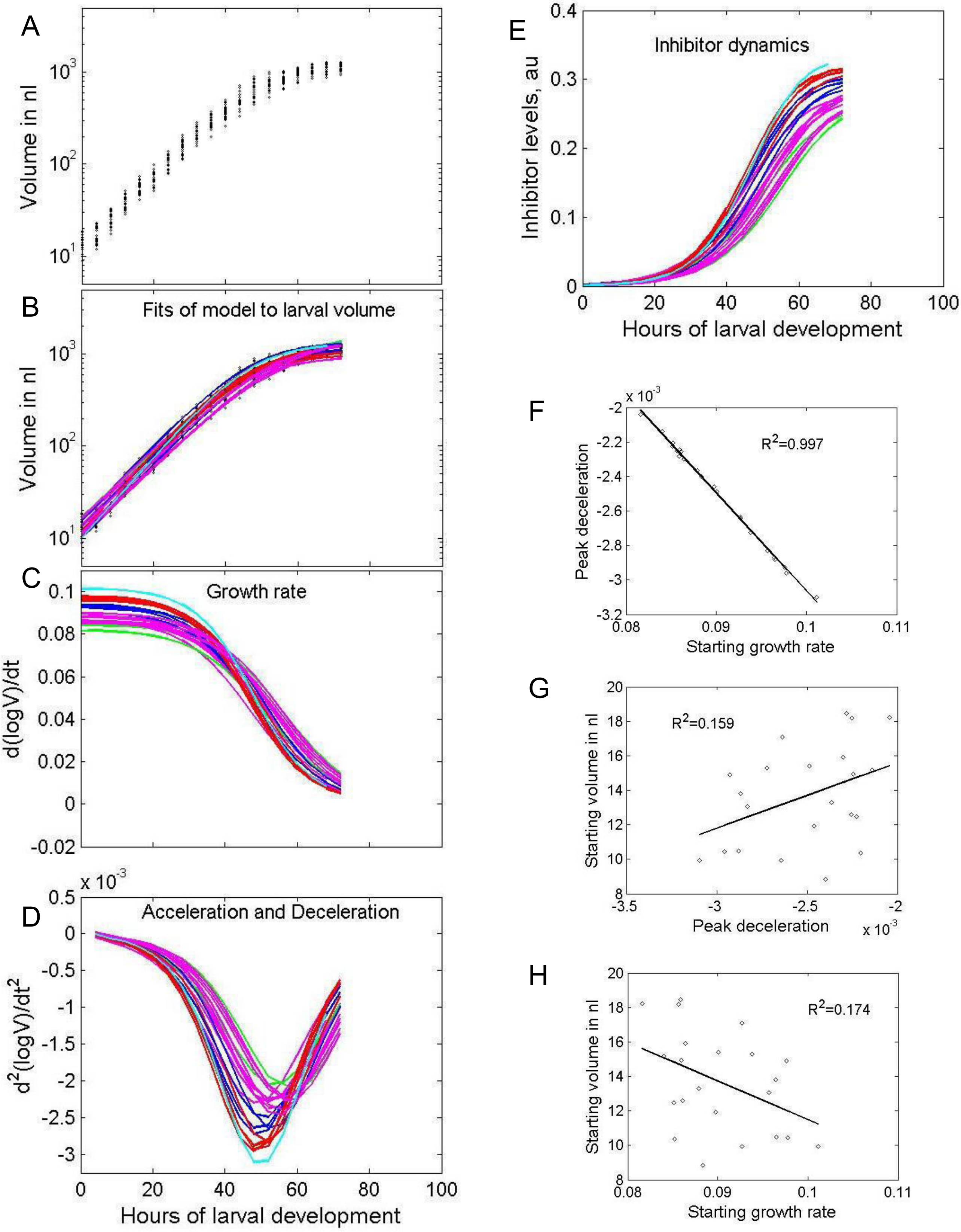


Supplemental Figure S7: Analysis of individual larval growth trajectories at 29°C
